# Close-Range Interactions Favor Growth in Random-Paired Extracted Soil Bacteria

**DOI:** 10.1101/646190

**Authors:** Manupriyam Dubey, Noushin Hadadi, Serge Pelet, David R. Johnson, Jan R. van der Meer

## Abstract

Species interactions at the cellular level are thought to govern the formation and functioning of microbial communities, but direct measurements of species interactions are difficult to perform between the hundreds of different species that constitute most microbial ecosystems. We developed a methodology to examine interactive growth of random cell pairs encapsulated inside 40–70 µm diameter agarose beads. We focused on a sandy soil as a test microbial ecosystem. By using gentle washing procedures, we detached microbial cells from sand and encapsulated them either in the absence or presence of pure culture inoculants. We then tested whether inoculants had on average positive or negative effects on the growth of resident community members depending on the growth substrate. Surprisingly, all the tested inoculants (including *Pseudomonas veronii* 1YdBTEX2, *Pseudomonas putida* F1, *Pseudomonas protegens* CHA0 and *Escherichia coli* MG1655) stimulated the growth of 40-80 percent of sand-derived cells when grown pair-wise in close proximity (i.e., within the same bead). This was true essentially irrespective of the growth substrate. Beneficial inoculant-sand cell partnerships resulted in up to 100-fold increase in productivity of the sand cell partner and up to 100-fold decrease in that of the inoculant. However, the maximum productivity attained by inoculant-sand cell partners within beads did not surpass that of inoculants alone. Further surprisingly, random pairs of sand cells encapsulated within the same bead also benefited growth in comparison to individual sand cells in a mutualistic manner (i.e., productivity when grown together was greater than the sum of individual productivities), but less than productivities observed in partnerships with the tested inoculants. This suggests that partnerships between inoculants and sand cells are not so much characterized by competition for substrate as by carbon loss through metabolite provision of the inoculant to sand cells (competitive exploitation).

## Introduction

Natural ‘free-living’ microbial communities and those in association with animal or plant hosts are exemplified by complex and high-density species interactions, being composed of dozens (e.g., certain insect hosts)^1^ up to many thousands (e.g., soils^2^) of individual species that live within short distances from each other (µm to mm–scale). Understanding the general principles of the formation, structure and functioning of microbial communities is one of the major questions in microbial ecology, and is still largely fragmentary^3-6^. Species compositions in communities vary, being subject to in- and outflow of species members^7^, to losses from selective predation^8,9^ or as a consequence of phage infection and lysis^10^, and as a result of fluctuating nutrient and chemical conditions in their environment. It is generally assumed that species interactions shape the community’s functioning within the physico-chemical boundary conditions of the system or the host^11-13^. However, given their complexity, species interactions within microbiota are challenging to dissect. Improved methodologies to infer interactions from community species characterization^11^, studies on simplified synthetic communities^14,15^ and mathematical modeling^11,16-18^ have helped to advance the understanding of community functioning^6,19-22^, but methods and studies that target complex and highly diverse systems are currently lacking^21^.

Frequent approaches to study microbial species interactions consist of coculturing two or three (labeled) species in spatially structured range-expansion experiments^23,24^, or inferring positive or negative interactions from species abundance fluctuations in defined well-mixed communities and conditions during prolonged culturing^15,25^. Our aim here was to develop a complementary approach that can assess species interactions from growth of randomly mixed cell pairs in small (ø40–70 µm) beads, as it had been suggested that pair-wise interactions are a reasonable predictor for interactions in higher order communities^15,26^. In particular, we aimed to study and infer the behaviour of pure cultures that can be used as inoculants to, for example, rationally manage, restore or complement existing communities with damaged functionalities (e.g., gut microbiome of diseased individuals, contaminated soils)^4,13,19,27^. Inoculants are typically selected for particular functional characteristics that are decisive for the intended complementation (e.g., expression of a xenometabolic pathway, expression of secondary metabolites for plant growth protection)^13^. On the other hand, there is only a limited knowledge base that could be used to predict survival and success of exogenously added strains within the target community, which may depend on the many interactions that the inoculant displays *vis-à-vis* the resident species members or vice-versa. Recent experimental studies have tried to infer inoculant behaviour by capturing genome-wide gene expression in complex environments like soil, but the number of detected differentially regulated functions has been too daunting to delineate simple complementation characteristics^28-30^. Our initial assumptions were thus that favorable inoculants may be differentiated by the magnitude of positive interactions with resident bacteria, but only on the type of substrate or condition they are functionally intended for.

To test this idea experimentally, we designed a methodology to randomly pair-wise encapsulate microbial cells within agarose beads, which are incubated in the presence of different growth substrates (Fig. 1, Table 1). Pair-wise productivity is then quantified from microcolony size estimations over time using microscopy imaging. As a test resident microbiota, we extracted and dispersed microbial cells from a sandy soil (*sand community* or SC). We then sought to encapsulate pairs of individuals into beads in the absence or presence of different inoculants (Table 1), with encapsulated inoculants serving as separate controls. Four inoculants were examined depending on their capacity to complement a xenometabolic pathway or because of their assumed competitive character. Xenometabolism was tested with *Pseudomonas veronii* 1YdBTEX2 (Pve)^30-32^ and with *Pseudomonas putida* F1 (Ppu)^33^, both capable of growing on toluene. We further tested *Pseudomonas protegens* CHA0 (Ppr), a plant growth promoting soil bacterium but which does not degrade toluene^34^, and *Escherichia coli* MG1655 (Eco), a non-native soil bacterium. In line with theory of microbial social interactions^35,36^, we expected that interactions between the many genotypes of soil bacteria would be dominated by competition (here: negative growth). Further, we expected that the inoculants Pve and Ppu would promote the growth of soil bacteria, but only in the presence of toluene. Namely, because toluene can have growth-inhibiting effects^37^, the consumption of toluene by Pve and Ppu should alleviate inhibition to soil cells and stimulate their growth. We assumed that the inoculant Ppr, on the basis of its capacity to produce antimicrobial secondary metabolites^38^, would yield on average less positive interactions, whereas Eco would be largely outcompeted in pair-wise growth with SC–cells. Surprisingly, however, all inoculants increased the productivity of sand-extracted bacteria and on any substrate, but only in close proximity interactions (i.e., within beads). Also close-proximity (within bead) presence of two or three SC–cells was on average positive on growth of either partner. In contrast, very little evidence for additive effects on productivity from inoculant addition was observed. Although demonstrated here for soil, our method is widely applicable for studying pair-wise growth interactions in any microbial community.

**Table 1.** Inoculant – sand community interactome experiments.

| Carbon Substrate | Inoculant |  |  |  |
| --- | --- | --- | --- | --- |
|  | Pve | Ppu | Ppr | EC |
| Mixed Carbon | +<sup>a</sup> | – | + | + |
| Sand Extract | + | – | – | – |
| Toluene | + | + | – | – |
| Succinate | + | – | – | – |

a) +, carried out ; –, not carried out.

**Fig. 1.**
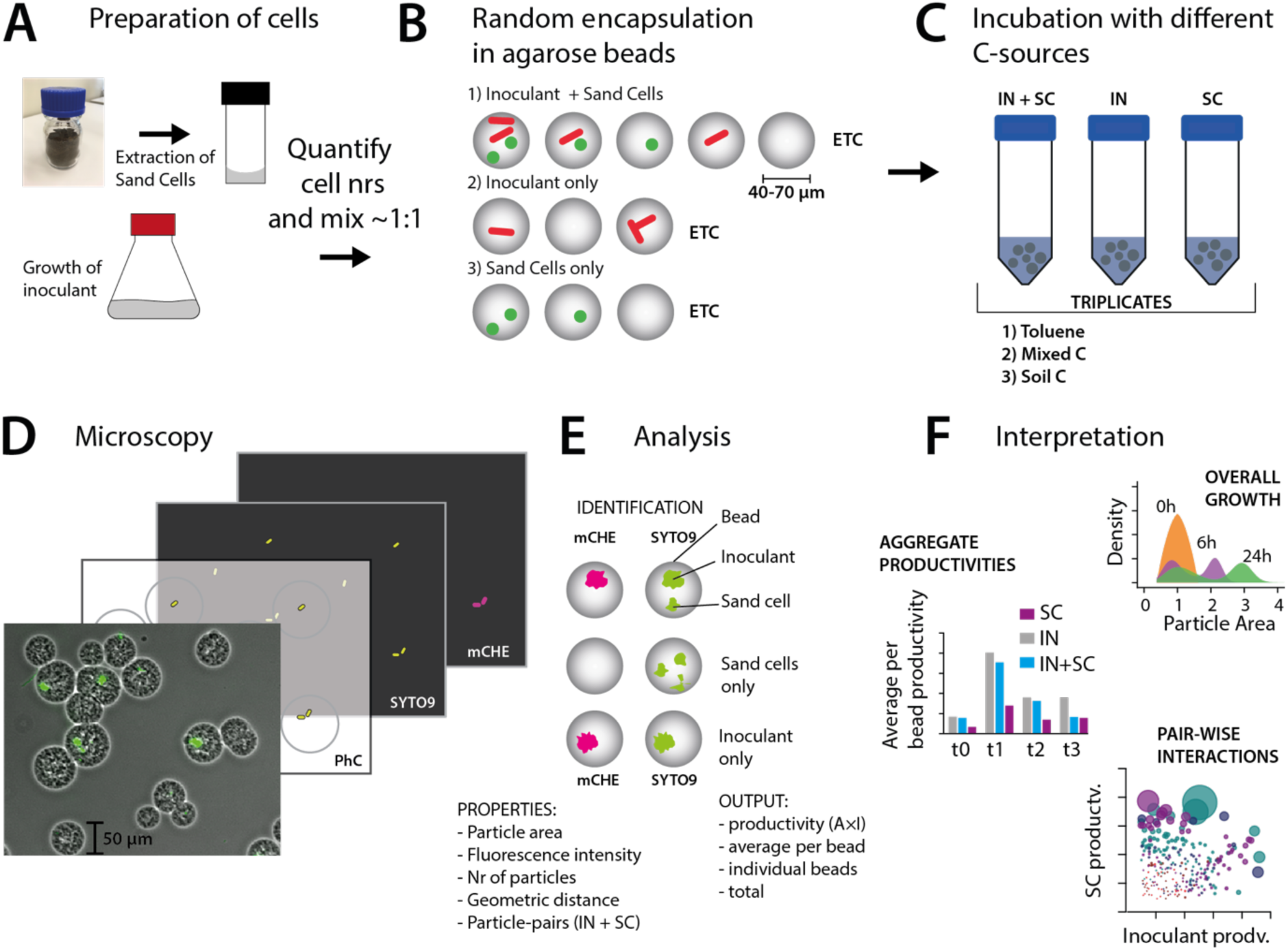
Workflow overview of the interactome approach. A. Sand community (SC) cells are extracted from fresh material and mixed with grown inoculant in equal ratio prior to encapsulation. B. Encapsulation generates random combinations of SC (green) and/or inoculant (red) in agarose beads having diameters between 40 to 70 µm. C. Beads of the three setups (IN, inoculant only; SC, sand cells only; IN+SC, inoculant plus sand cells) are suspended and incubated in relevant growth media, and are sampled over time. D. Bead microscopy at five timepoints, and 10 random image positions, imaging in phase contrast (PhC), mCherry fluorescence (mCHE) and SYTO-9 fluorescence. E. Data analysis pipeline identifies beads, and measures properties of microcolonies of inoculant (mCHE+SYTO-9) and sand cells. F. Output data types at system’s or pair-wise interaction level.

## RESULTS

### Encapsulation and growth measurements of randomized pair-wise strain combinations

To develop a system to infer potential species interactions in a complex microbial community we studied randomized pair-wise growth of dispersed community-derived cells, without or with intermixed specific bacterial inoculants, and on a variety of externally added substrates (Fig. 1, Table 1). We chose agarose microbeads as containers to impose long-term close spatial proximity (<70 µm) of the starting cells, assuming that if their interaction would be positive, the cells would divide and form detectable microcolonies, whereas if it would be negative, one or both of the starter cells would not or poorly divide. Long distance interactions (i.e., between beads) may also occur, as agarose beads are permissive for diffusion of small molecules (e.g, substrates, metabolic intermediates), but we expect their influence to be minor given the dilution in the experimental setup. Growth of cells over time was inferred from the changes in biovolume of the microcolonies within individual beads, estimated from the fluorescence area taken up by SYTO9-stained cells times their mean fluorescence intensity (to account for multiple cell layers). Inoculants were differentiated specifically from resident community species by genetic labeling with a constitutively-expressed mCherry fluorescence protein (Fig. 1). As the starting composition of cells in the beads was Poisson random (aiming at 1–2 cells per bead), the image-analyzed bead mixtures contained both single, double or higher numbers of microcolonies, of SC–cells alone or in combination with one of the inoculants (Fig. 1).

We tested four different inoculants (*P. veronii* [Pve], *P. protegens* [Ppr], *P. putida* [Ppu] and *E. coli* [Eco]) and four different carbon substrate regimes (succinate, toluene, a mixture of 16 C-substrates, and a ‘sand extract’; although not in all combinations, Table 1). When grown alone, all the inoculants were able to divide and form colonies inside agarose beads, given that cell microcolony sizes increased over time of the incubation in the different deployed growth media (Fig. 2, Fig. S1). The size of Pve and Eco cells at time of inoculation in the beads was slightly larger than that of Ppr and Ppu and that of a (typical) SC–cell after extraction from soil (Fig. S1). To estimate the maximum number of cell divisions, we compared the maximum microcolony area range after 24–72 h of growth for the inoculant compared to that at time of inoculation (Fig. 2). Assuming a round microcolony with densely packed cells, the observed increase of 50–100-fold would correspond to maximum cell numbers of approximately 450–1000 from a single starting Pve or Ppr cell (∼9–10 generations). Importantly, our method thus enabled us to isolate ecological processes (i.e., species interactions) from evolutionary processes that occur over longer time-scales, such as the appearance of genetic changes that could modulate growth properties. Microcolonies from encapsulated cells from the sand community also increased in size over time (Fig. 2, SC), confirming their growth, albeit to a smaller extent than for the inoculants. If we assume the decrease in the proportion of the smallest microcolony area compared to time=0 as being indicative for the proportion of SC–cells capable of dividing in the respective medium, we estimated some 20 % of SC–cells to divide within 6–24 h (Table S1, depending on the substrate). Microcolony size distributions in the coculture incubations (e.g., inoculant plus SC) were on first sight a combination of that of inoculant and SC incubated separately, with characteristic ‘peaks’ appearing during incubation, attributable to, e.g., Pve or Ppr (Fig. 2A, B), suggesting inoculant cells to grow in coculture with SC.

**Fig. 2.**
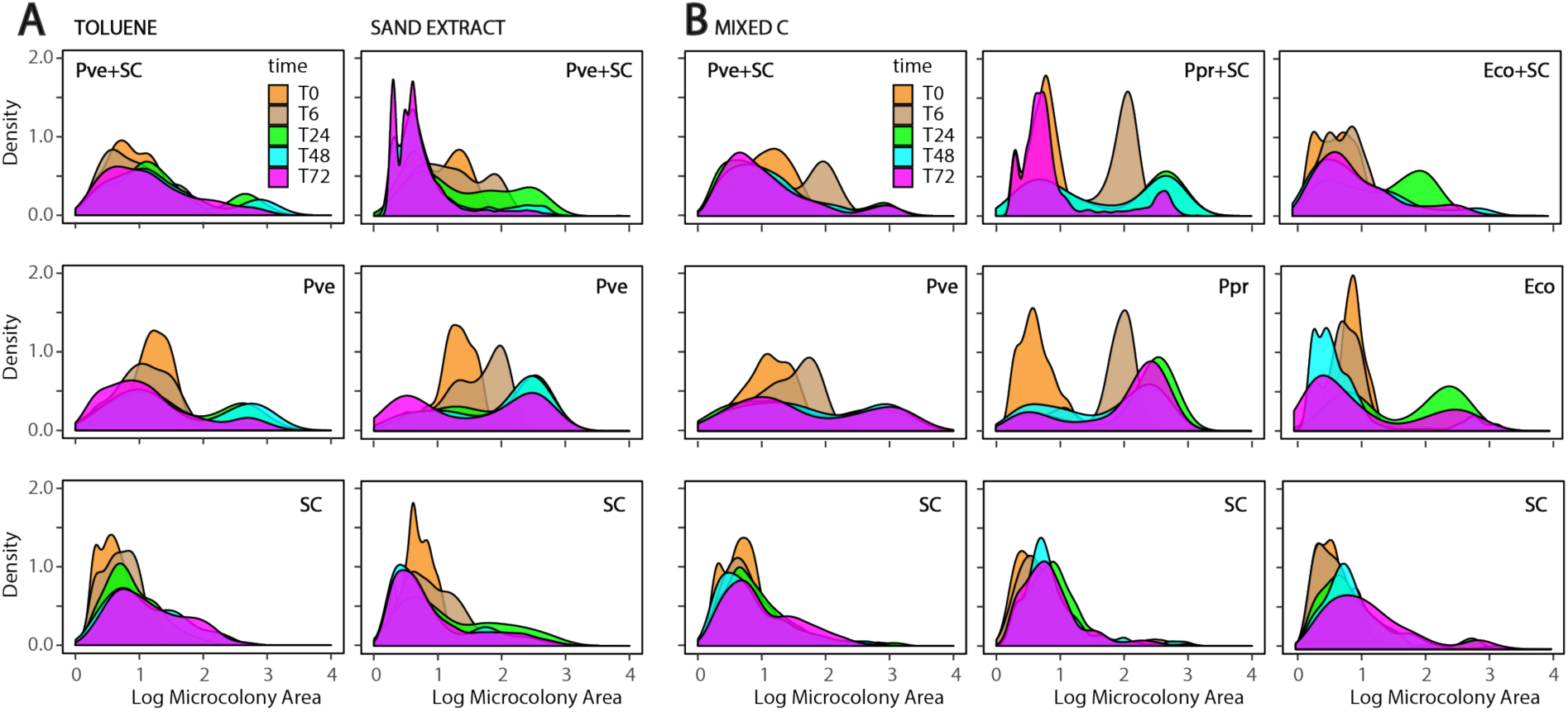
Changes in observed microcolony sizes over time as function of inoculant and growth substrate. A. Microcolony sizes of *P. veronii* (Pve) in presence or absence of sand community (SC) cells grown with toluene or sand-extract. B. Microcolony sizes of three different inoculants (*P. veronii,* Pve; *P. protegens,* Ppr; *E. coli*, Eco) in presence or absence of SC grown in mixed-C substrate. Microcolony sizes represented as the ^10^log measured area. Density, the Gaussian frequency distribution of the microcolony areas. Sampling time points (T0, T6, etc.) indicated in h.

### Aggregate productivity in the presence or absence of inoculant

In order to compare the total productivities across the various incubations as aggregate properties of all beads, but considering that the assays had varying amounts of beads, we normalized the observed biomass growth (as particle area times fluorescence intensity) on a per-bead basis for each experiment. As an example, Figure 3 summarizes per-bead productivities of incubations of SC–cells without or with the inoculant Pve on different carbon substrates. The normalized per-bead productivity of Pve (Fig. 3, magenta bars) surpassed that of SC-cells (Fig. 3, cyan bars) on all four carbon substrates (p<0.001, ANOVA followed by post-hoc Tukey test). The type of substrate affected the observed productivities, but since the total amount of C in the case of ‘sand extract’ and toluene may be different than for mixed-C or succinate (both at 0.1 mM C), we cannot test for this significance.

**Fig. 3.**
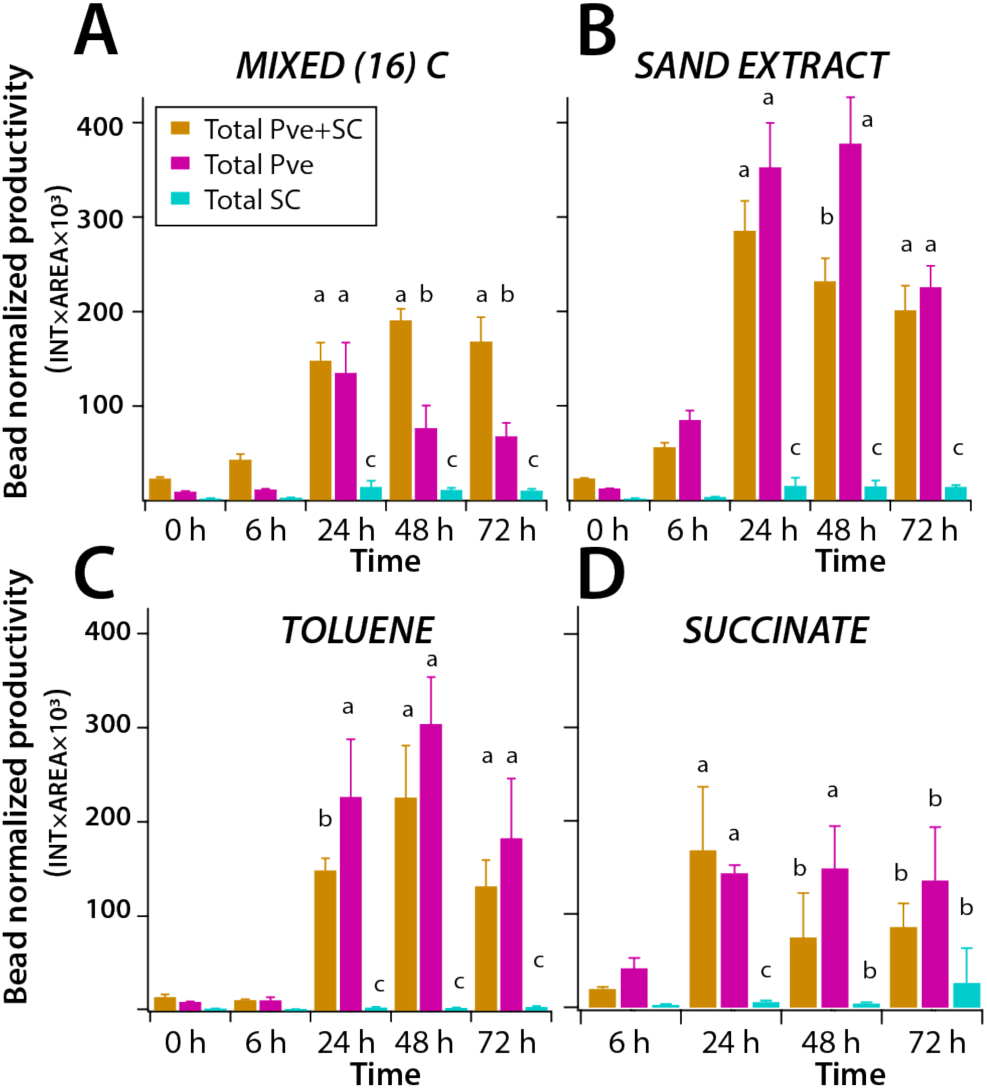
Global productivity of sand community cells in presence or absence of *P. veronii* as a function of time and growth substrates. A. Mixture of 16-carbon substrates (0.1 mM total). B. Sand extract solution. C. Toluene (provided through the gas phase). D. Succinate (0.1 mM). Bars show mean productivity, defined as the product of particle area times particle fluorescence intensity, normalized across all observed beads per series time points. Error bars represent the standard deviation of the mean in triplicate incubations. Letters indicate significance groups in ANOVA followed by post-hoc Tukey testing (a–b, p<0.01; a–c, p<0.001).

Although different in individual experiments and at singular time points (e.g., Fig. 3), the maximum normalized per-bead productivity at any time point of the coculture incubations (Pve+SC) on any of the substrates was statistically significantly smaller from that of Pve alone, across multiple independently repeated experiments (Fig. S2, p=0.04614, Wilcoxon rank sum test on medians). This suggests that growth of Pve–cells at system’s level (i.e., the incubation vial) is negatively influenced by the presence of SC–cells, resulting in a smaller yield of the inoculant. Across all incubations, the maximum per-bead productivity of the SC incubations was less than either Pve or Pve+SC (Fig. S2, p<0.001, ANOVA followed by post-hoc Tukey test). Importantly, the difference in mean per-bead productivity between Pve and Pve+SC was statistically the same as the productivity of SC alone (p=0.7244, Wilcoxon rank sum test). This suggests there were on average no additive effects and total productivity was determined by the total available carbon substrate. The per-bead productivity increased statistically significantly in two out of five Pve+SC cocultures compared to Pve alone, in case of the mixed-C substrates (Fig. 3A, Fig. S2, p=6.84×10^−5^ and p=0.0055 in paired t-test). For those particular incubations, one might conclude that the growth interactions between partners at system’s level had been positive and additive (i.e., yielding higher biomass than either achieved in separation). The reason for this may be that the sand community was extracted freshly at each occasion and assay from samples taken at the natural location, and may have constituted a slightly different starting species composition.

### Inoculant- and substrate-dependent system productivity

The normalized per-bead productivity averaged across the assay as a whole did not illustrate specifically the types (e.g., positive or negative) or extent of interactions between inoculant and SC–cells occurring inside the same beads. The reason is that the encapsulation process is random and Poisson-distributed. Therefore, even though the inoculant (e.g., Pve) is mixed with SC–cells in the agarose, they do not necessarily end up within the same bead (as schematically illustrated in Fig. 1B). To get a better picture on the average interactions when inoculant and SC–cells grow inside the same bead, we discriminated in the coculture (e.g., Pve plus SC) for beads that carried (by chance) only Pve colonies, for only SC colonies, and for those that carried both Pve and SC colonies. Their mean productivity was subsequently compared to incubations on the same carbon substrate(s) with Pve or SC–cells separately. On mixed-C substrates this indicated, for example, that Pve incubated separately without SC (Fig. 4A, mixed carbon, dark green bars) had a far greater productivity than either Pve alone in beads but combined with SC–cells in the same flask (Fig. 4A, green bars), or Pve within the same beads as SC–cells (orange bars, ANOVA with post-hoc Tukey test, p<0.0001). However, the latter two did not have statistically significantly different productivity. In contrast, SC–cells in the same experiment had a far greater productivity when they found themselves within the same beads as Pve (p<0.005, ANOVA with post-hoc Tukey test, Fig. 4B, mixed carbon). This explains the average loss of Pve productivity in the same beads, but it is curious that both beads with Pve or SC alone that occur in the mixed Pve+SC incubation did not profit from a significant increase in productivity. A similar situation occurred on sand extract as carbon regime (Fig. 4). Pve incubated separately on sand extract had a higher per-bead productivity than in combination with SC–cells (p<0.005, Fig. 4A), but in this case Pve–cells that happened to be co-encapsulated with SC–cells within the same beads on average fared better than Pve–cells that occurred alone in beads in the mixed incubation (Fig. 4A, SAND EXTRACT, p<0.05). SC–cells again showed the highest per-bead productivity when they found themselves with Pve inside the same bead (p<0.005), but only slightly significantly lower when they occurred alone inside the combined incubation (Pve + SC, p<0.05, ANOVA followed by post-hoc Tukey test, Fig. 4B). This suggests that under those conditions, SC– and Pve–cells on average interacted positively within the close range of the same bead, which increased productivity. In the extreme case of toluene as substrate, which only Pve and Ppu can metabolize, the Pve per-bead productivity declined both in the mixture with SC and even more inside the same beads as SC (Fig. 4A, TOLUENE, p<0.001), suggesting that part of the toluene metabolites produced by Pve were being lost and utilized by SC–cells. On the other hand, the mean SC productivity under those conditions was very small and not significantly different for SC–cells within the same beads as Pve (Fig. 4B, ANOVA followed by post-hoc Tukey test).

**Fig. 4.**
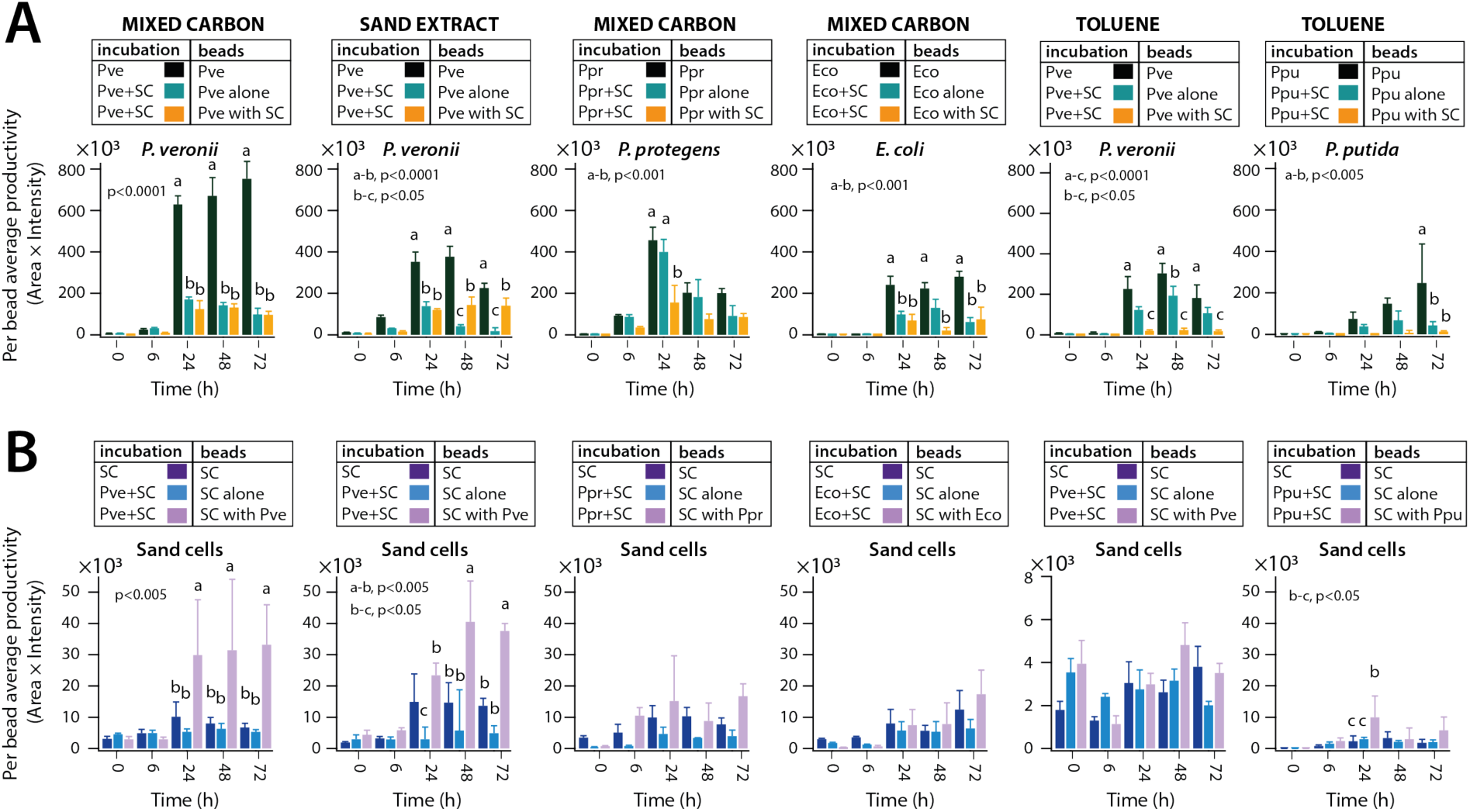
Productivities of inoculant in presence or absence of sand cells within the same bead. A. Productivity for inoculant incubation alone (e.g., *incubation* Pve, *beads* Pve), or for incubation in presence of sand community cells (Pve+SC), either for beads with Pve alone, or for beads with Pve and SC co-occurring. B. As A, but for productivity of sand community cells. Bars indicate the mean per–bead productivity across triplicate incubations. Error bars denote standard deviations from the mean. Small letters indicate significance levels in ANOVA across time series and categories followed by post hoc Tukey testing. Plots in A and B below each other correspond to the same incubation and carbon substrate.

To place these observations with Pve as inoculant in perspective, we repeated experiments on mixed-C substrates with Ppr (as an assumed competitive soil bacterium) and with Eco (as atypical soil bacterium), and on toluene with Ppu (as another toluene degrading bacterium). In contrast to Pve, Ppr displayed the same mean per-bead productivity whether incubated alone or finding itself alone inside agarose beads in the mixture with SC (Fig. 4A, Ppr, ANOVA followed by post-hoc Tukey test). Also the mean productivity of SC–cells was not statistically significantly different among beads with a direct Ppr partner, or alone in incubation (Fig. 4B, ANOVA). Eco productivity on mixed-C substrates was clearly lower than that of Ppr or Pve, and was lower in combination with SC–cells, irrespective of being inside the same beads or alone in beads in the mixture (Fig. 4A, Eco, p<0.001). In contrast to Pve, the mean productivity of SC–cells did not significantly increase in combinations with Eco (Fig. 4B, Eco). In comparison to Pve on toluene, Ppu took slightly longer to develop the same mean productivity and Ppu–cells incubated with SC were less productive (Fig. 4, TOLUENE, p<0.005 t=72 h, ANOVA followed by post-hoc Tukey). In contrast, SC–cells in the same beads with Ppu hardly increased their productivity than in its absence (Fig. 4B, TOLUENE, p<0.05, t=24 h, ANOVA). In terms of aggregate productivity these results suggested, therefore, that SC–cells develop better with Pve than with the other inoculants, irrespective of the growth substrate (Fig. 4B).

### Pair-wise growth analysis suggests widespread positive effects of inoculants

Even though, in some cases the mean per-bead productivity of SC–cells increased in presence of inoculant cells within the same bead (e.g., Fig. 4), the process of averaging masked the types and extent of individual pair-wise interactions. Next, therefore, we analysed solely those beads having at least one inoculant and at least one SC microcolony. We scored the individual microcolony sizes of both partners, as well as those observed in individual beads of either SC– or inoculant cells incubated separately (Fig. 5). As example, in the case of toluene as substrate, SC–cells alone on average developed very poorly (as displayed in Figs 3 and 4), but those SC–cells being within beads with Pve developed much more strongly. Now, whereas the mean per-bead normalized productivities were not statistically significantly different for SC growth in presence or absence of Pve (e.g., Fig. 4B, sand cells, TOLUENE), individual bead analysis indicated that 42.6% of all beads with Pve partners at any time point led to SC growth above the 95th percentile of SC alone (Fig. 5A, Table 2, p=1.80×10^−5^), which corresponds to a 100-fold increased productivity of SC (Fig. S3). In contrast, the productivity of those Pve–cells in beads together with SC–cells decreased by 100-fold, and in no single case Pve–cells profited from SC–cells in their productivity compared to the 95^th^ percentile of its growth alone (Fig. 5A, Table 2, p=5.4×10^−5^). The distribution of summed productivities of SC+Pve among those beads in which SC-cells largely profited (>95^th^ percentile) was statistically significantly different from that of Pve cell productivities incubated on toluene separately (Fig. S4, p=0.005 Fisher’s exact test for productivity distributions). This suggests that the productivity of SC+Pve incubations on toluene did not surpass that of Pve alone.

**Table 2.** Productivity increase or loss in inoculant– sand community (SC) cell pairs compared to either inoculant or SC alone.

| Carbon Substrate | Bead Combination | Productivity increase<sup>a</sup> |  |  |  | Percentage no growth<sup>b</sup> |  |  |  |
| --- | --- | --- | --- | --- | --- | --- | --- | --- | --- |
|  |  | Inoculant | SC | Inoculant & SC | p-value<sup>c</sup> | Inoculant | SC | Inoculant & SC | p-value |
| Mixed Carbon | Pve+SC | 0 | 46.6 | 0.3 |  | 8.5 | 8.2 | <0.1 |  |
| | Pve<sup>d</sup> | 2.7 | | | $4.03 \cdot 10^{-7}$ | 9.2 | | | 0.4345 |
| | SC<sup>d</sup> | | 7.6 | | $1.21 \cdot 10^{-12}$ | | 7.4 | | 0.2832 |
| Mixed Carbon | Eco+SC | 0.4 | 48.4 | 0.8 |  | 11.9 | 2.3 | 0 |  |
| | Eco | 8.1 | | | $7.10 \cdot 10^{-5}$ | 15.1 | | | 0.4632 |
| | SC | | 7.3 | | $4.30 \cdot 10^{-5}$ | | 9.2 | | 0.0138 |
| Mixed Carbon | Ppr+SC | 0.1 | 80.1 | 0.5 |  | 10 | 2.5 | <0.1 |  |
| | Ppr | 1.2 | | | $4.10 \cdot 10^{-7}$ | 1.6 | | | 0.0021 |
| | SC | | 4.8 | | $1.20 \cdot 10^{-12}$ | | 7.3 | | 0.0103 |
| Sand extract | Pve+SC | 0.2 | 44.5 | 0.4 |  | 4.5 | 5.8 | 0 |  |
| | Pve | 2.4 | | | $2.60 \cdot 10^{-6}$ | 1.8 | | | 0.0258 |
| | SC | | 3.8 | | $9.90 \cdot 10^{-13}$ | | 12.1 | | 0.0332 |
| Toluene | Ppu+SC | 0.9 | 27.2 | 0.4 |  | 10 | 22.7 | <0.1 |  |
| | Ppu | 10.2 | | | $5.40 \cdot 10^{-5}$ | 12.7 | | | 0.2085 |
| | SC | | 5.6 | | $1.80 \cdot 10^{-5}$ | | 11.1 | | 0.0904 |
| Toluene | Pve+SC | 0 | 42.6 | 0 |  | 4.3 | 11.9 | 0 |  |
| | Pve | 3.2 | | | $7.90 \cdot 10^{-16}$ | 0.7 | | | 0.0144 |
| | SC | | 5.6 | | $1.30 \cdot 10^{-7}$ | | 5.7 | | 0.0079 |

d) Inoculant or SC incubations separately on the same substrate.

**Fig. 5.**
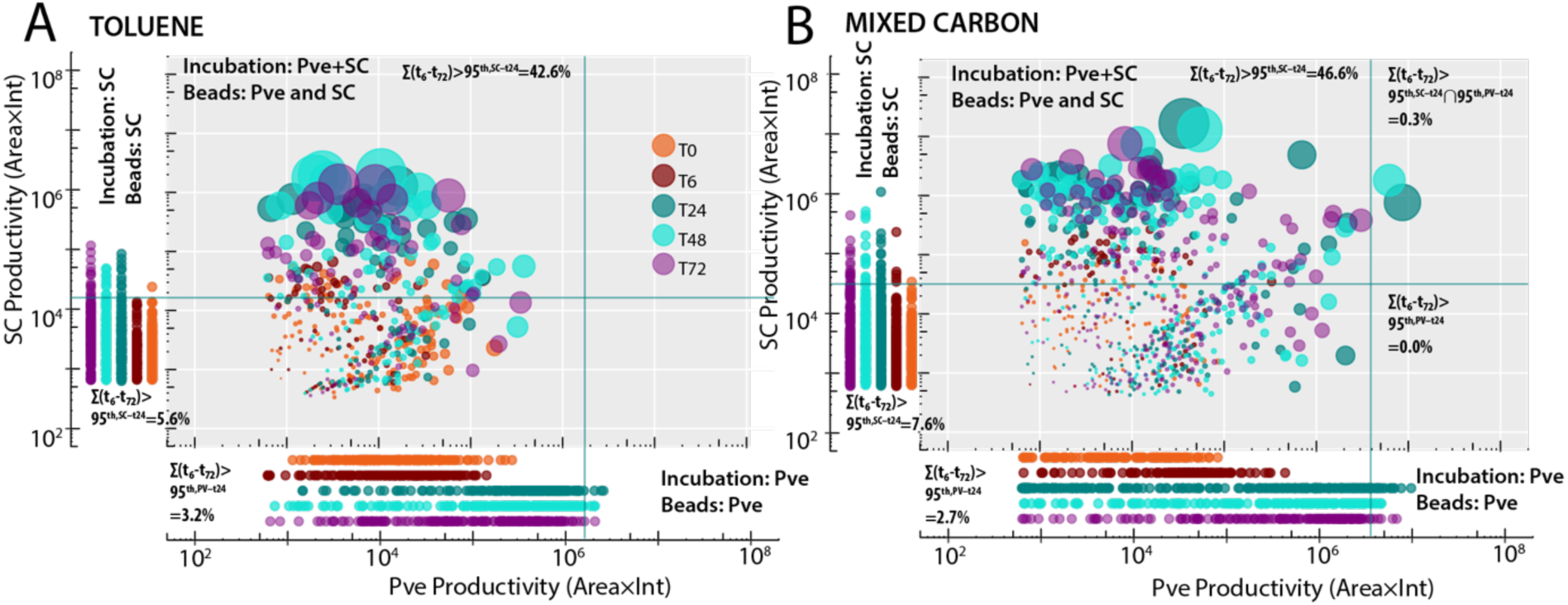
Pair-wise productivities within beads containing both inoculant and sand community cells. A. Individual pair-wise productivities on toluene of co-occurring *P. veronii* (Pve) and sand community (SC) within the same bead (colored bubbles), compared to productivity of Pve (bottom) or SC (left side) in separate individual incubations. B. As A, but for mixed-C substrates. Bead productivities are displayed on log-axes. Green lines indicate the 95^th^ percentile productivity of the individual incubation at t=24 h. Bubble diameters represent the Euclidian distance from the origin and are a relative measure of the microcolony sizes. Percentages indicate the proportion of beads of the total, falling above the respective 95^th^ percentile threshold (values summarized and tested for significance in Table 2).

In the incubation with mixed-C substrates the general productivity by SC–cells was much higher than on toluene, but compared to the 95^th^ percentile at 24 h a further significant increase was observed for 46.6% of the beads with Pve partnerships at any time point (Fig. 5B, Table 2, p=1.21×10^−12^). This indicated that despite containing a mixture of very general carbon substrates, half of the SC–cells at start still profited from being in the same bead with a Pve inoculant cell. Interestingly, on mixed–C substrates we also observed a small percentage of beads where both SC and Pve profited of being together (Fig. 5B). With sand extract as substrate, SC–cells on average had the highest productivity, but in coculture still 44.5% of beads profited from partnering with Pve (i.e., >95^th^ percentile of SC–cells incubated separately, Table 2, p=9.9×10^−13^), and with 0.4% of beads where both partners profited (Fig. S5, Table 2). For this subclass of SC+Pve beads (with SC>95^th^ percentile) the distribution of summed productivities was shifted to higher values than that of Pve alone (Fig. S4, p=0.0005 in Fisher’s exact test).

In case of Ppr as inoculant and with mixed–C substrates, SC profited even more (80.1% of beads with increased growth, and 0.5% with increased growth of both SC and Ppr; Fig. S5, Table 2, p=1.2×10^−12^), but without significantly increased productivity of this subset of cocultured beads compared to Ppr growing on mixed-C alone (Fig. S4, p=0.0645). Eco as inoculant with mixed–C substrates yielded similar proportional benefits to SC–cells as Pve, i.e., 48.4% (Fig. S5, Table 2, p=4.3×10^−5^). A further small 0.8% proportion of beads occurred in which both SC and Eco had profited (Table 2), and, interestingly, the distribution of summed productivities for SC+Eco partnering beads was significantly different from that of Eco alone, suggesting that the inoculant profited to some extent from being with SC–cells (Fig. S4, p=0.0005 in Fisher’s exact test). Ppu as inoculant was profitable to partner SC–cells on toluene in 27.2% of beads (Fig. S5, Table 2, p=1.8×10^−5^). The distribution of summed productivities for Ppu+SC pairs was statistically significantly shifted to lower values than that of Pve+SC pairs (Fig. S4, p=0.0005 Fisher’s exact test), suggesting SC–cells profit less from Ppu than from Pve as inoculant on toluene.

In order to infer the magnitude of potential negative interactions, we estimated the loss in productivity of the inoculant in pairs with SC–starting cells, and the percentage of inoculant–SC pairs under the different conditions where one of the partners did not grow at all (Table 2). In comparison to the inoculant incubations alone, all inoculants significantly lost productivity when in the same bead with SC–cells (Table 2, p-values between 10^−5^ and 10^−17^ for the decrease of boosted average inoculant microcolony sizes). In comparison to either partner alone on the same substrate, in case of general substrates a lower percentage of SC–cells (2.3–8.2%) did not seem to grow at all in partnership (i.e., particle area × intensity at 24, 48 or 72 h < 10^th^ percentile at T=0, Table 2, p=0.0103–0.0332), and in less than 0.1% of all pair-wise combinations no growth of either partner occurred. Notable exceptions were for growth on toluene, where the percentage of non-growing SC–cells with Ppu increased to 22.7% and with Pve to 11.9% (Table 2, p=0.0904, p=0.0079, respectively). In contrast, in three cases the percentage of non-growing inoculant slightly increased in combination with SC–cells to between 4.3–11.9% (Table 2, p=0.0021– 0.0258). This suggested that maximally some 10% of pair-wise interactions might be inhibitory on one of the partners.

### Interactions between sand cell partners are positive for productivity

Finally, the bead interactomes also contained numerous cases of only SC–SC partnership beads that randomly contained two or more starting SC–cells (without any inoculant). When pooling all such beads of SC–SC partnerships from the experiments conducted either in mixed–C or sand-extract as growth substrates, and ordering the biggest partner on the x– and the smaller partner on the y–axis, we could see that around two-thirds of SC-SC partnerships are dominated by one big and one small microcolony (i.e., more than two-fold size difference). In about one-third of cases, both SC microcolonies inside single beads are less than two-fold different (Fig. 6A, B). No effect of the distance between both SC-microcolonies on their mutual size was discernable (Fig. 6C, r^2^=0.00032), which might have been intuitively expected. In contrast, for SC–cells in pairs within the same bead (without inoculant), and across both substrates, there was an overall strong overproportional increase (i.e., more than twice) of per–bead productivity compared to that of single SC–microcolonies (Fig. 6D, p<0.05 for t=24 h, p<0.005 for t=48 or 72 h). There was no further additive effect when beads contained, by chance, more than two SC–microcolonies (Fig. 6D, yellow box-plot data, p=0.3229–0.7611). This suggests that, even though in the majority of cases, paired SC–cells grow to unbalanced microcolony sizes (i.e., more than twofold different), their interactions are on average positive for their collective productivity.

**Fig. 6.**
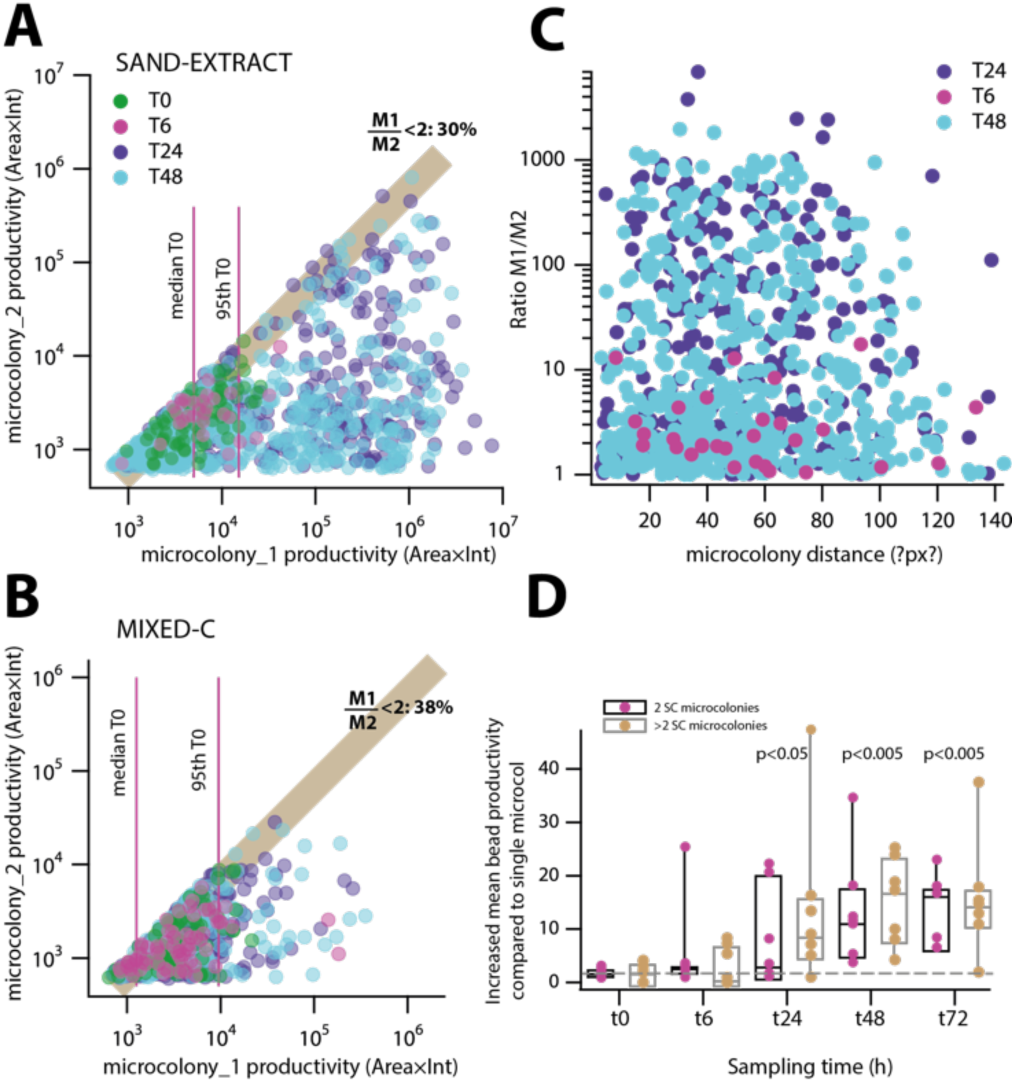
Pair-wise interactions among sand community members. A, B. Grouped beads with exactly only two SC–microcolonies under sand-extract (A) or mixed carbon substrate (B) regime. Pairs are ordered arbitrarily with the largest microcolony on the x– and the smaller microcolony on the y–axis. Bead colors indicate sampling time points (in h). The light brown shading and corresponding percentage indicate the proportion of pairs with a productivity difference of less than 2 (ratio M1/M2 <2). Cyan lines denote the median and the 95^th^ percentile microcolony productivities of all SC–pair beads at t=0. C. Paired microcolony ratio (M1/M2) as a function of geometric distance of the microcolony centres. D. Mean bead productivity increase in SC–pairs or higher SC–microcolony numbers compared to single SC occupancy across all incubations and substrates. Data points represent the means from the individual incubations and substrates, inside box plots. The dotted line denotes the expected increase in productivity of an additive SC–pair compared to single occupancy at t=0 (i.e., twofold). P-values are derived from pair-wise t-test comparisons of the productivity measurements in all seven experimental conditions for beads with two or more than two SC members, to that of single occupancy beads at the same sampling time point.

## DISCUSSION

We developed a method to measure growth interactions from complex mixtures of microbial cells randomly encapsulated inside individual agarose beads. We tested our method on resident microbes washed from a sand-microbiota system, and incubated in absence or presence of specific inoculant bacteria, in order to compare whether inoculants may be positive for growth of the resident community. Our method permitted to follow pair-wise growth in hundreds of beads simultaneously that can be averaged to assess mean normalized bead productivity changes as a function of carbon substrate(s) (e.g., Fig. 3) or inoculant type (Fig. 4). The method further provides estimates of the proportions of positive and negative growth interactions within the resident community members itself, or in presence of inoculant (e.g., Fig. 5 and 6, Table 2).

Most strikingly, the sand community as a whole (Fig. 4B) and a substantial subset of SC–cells in particular (27-80%, depending on inoculant and substrate, Fig. 5, Fig. S5, Table 2), increased productivity in presence of the inoculant, but only in case of close proximity of growing within the same bead. In contrast, productivity of all inoculants under all conditions declined in pair-wise growth with SC–cells in close proximity (i.e., being within the same bead, Fig. 5, Table 2), but mostly also at larger distance (i.e., inoculant alone in a bead, but in the same mixture with encapsulated SC–cells, Fig. 4). On average, in some 10% of the inoculant–SC pairs one of the partners did not develop (Table 2), suggesting clear negative (inhibitory) interactions. Finally, these effects occurred almost irrespectively of the type of inoculant, although some inoculants (notably Ppr) yielded extremely positive effects on SC–cells (80% above the 95^th^ percentile of SC growth alone). Mostly, the summed productivities of beads with partnering inoculant and SC–cells did not surpass that of inoculants on the same substrate alone (Fig. S4). From these results we thus conclude that the used inoculants were in close proximity interactions beneficial for the productivity of dispersed resident cells from the sandy soil.

Our results suggest that the majority of interactions between inoculants and SC–cells in close proximity are highly imbalanced, with SC–cells increasing their mean productivity by a factor of 100 and inoculants losing productivity by a factor of 100 (Fig. 5, Fig. S3). These imbalanced interactions are neutral with respect to total productivity (i.e., distribution of summed SC+inoculant bead productivities does not surpass that of inoculant alone, Fig. S4), but the SC–cells are largely dependent on the inoculants, since without their presence the SC productivity is very low. This suggests that it is not so much competition for growth substrate per se that dominates these interactions, but either specific growth factors (e.g., amino acids) or metabolic intermediates (e.g., acetate) produced by the inoculants that favor excessive growth of a large proportion of SC–cells. Instead of competition (both partners suffer), therefore, this type of interactions might be defined as competitive exploitation (rather than parasitism^11^). In a small proportion of cases (∼0.5%, at >95^th^ percentile of individual growth, Table 2), and pronounced on sand extract as substrate (Fig. S4), we found evidence for additive effects on productivity of both inoculant and SC–cells. If one would consider the additive case as being representative for cooperative behaviour, it would confirm the theory that only a minority of social interactions in communities is cooperative^36^. However, results from beads of co-occurring SC–SC pairs within the same experiment (Fig. 6) suggest that also among the sand community itself, it is largely profitable for productivity to be with partners (although this productivity increase does not rival the increase obtained from being paired with any of the inoculants). This result is therefore a strong indication that partnerships within close distance in complex communities are favorable for increased productivity, perhaps as a result of metabolite exchange in general or metabolites favoring growth of auxotrophs^39,40^. The inferred behaviour of SC–inoculant partnerships (i.e., competitive exploitation) may thus not be representative for the majority of interactions within complex carbon-limited communities as in soil that show higher-than-additive productivities (i.e., mutualistic interactions) in pair-wise random incubations in close proximity (Fig. 6D).

One of our initial assumptions was that inoculants would specifically favor SC productivity in the case of xenometabolic complementation (here: toluene) and less so for general available substrates. SC productivity was overall impaired on toluene in comparison to other substrates (Fig. 4B). Although we did find a strong positive effect of the two toluene-degrading inoculants Pve and Ppu on SC growth in pair-wise analysis (Fig. 5A), this was not very well visible in the mean bead productivities (Fig. 4B), and the proportion of non-growing SC–cells even slightly increased (Table 2), suggesting that the beneficial effects may be limited as a result of the nature of the specific metabolites which are leaking from toluene-metabolizing cells (e.g., catechols). To our surprise, however, the inoculants were also extremely beneficial for SC–productivity on the other types of carbon substrates (Fig. 5B, Fig. S5, Table 2). Even more surprising was that Ppr, considered to be a strong competitor^34^, actually provided the largest measured benefit on SC–cells, with 80% of observed pair-wise SC+Ppr interactions having SC productivities above the 95^th^ percentile of SC growth alone (Table 2). The major benefit of all inoculants on the sand community could thus be their capacity to provide growth factors or direct central metabolites from the primary carbon substrates to community members, which is reminescent of the types of carbon sharing reported in simplified communities^41^. When assuming that the proportion of beneficial pair-wise interactions on SC cells (>95^th^ percentile of its own incubation) is an indication for the potential ‘hub’ of interactions in a community network^5,42^, one might consider that a species like Ppr is a strong potential keystone member for a soil community (in case it is sufficiently abundant).

Productivity measurements to assess strain-strain interactions have typically been based on cell density changes in suspended growth^15,25^, on species abundance fluctuations in natural-derived communities^41^, or on expansion-range experiments from droplet cocultures placed on agar plates^23,24^. Both of these cannot readily be parallelized for multiple strain-strain interactions. The agarose bead containers deployed in our experiments supported microcolony growth at the expense of diffusing carbon and nutrients from the medium, and allowed simultaneous growth monitoring of hundreds of pairs. Pair-wise growth interactions were shown to be reasonably good predictors for septet and octet artificial communities^15^, and thus, an upscaling of observable pair-wise interactions as demonstrated here, may help to extrapolate functional behaviour in more complex communities. Our method is widely amenable to different starting communities, given the ease of bead encapsulation. The methodology can be adapted by limiting long-distance interactions as a result of nutrient- and metabolite exchange to the medium, through the use of bead-in-oil emulsions or other type of bead variations. The method may be further improved by combining bead growth analysis to either a global analysis of the (changes) in species compositions of the targeted community in the bead incubations, or to methods that would identify species pairs on a per-bead basis^43,44^. In conclusion, the randomized pair-wise bead encapsulation growth methodology is widely applicable to study the collective growth interactions (the *interactome*) of microbiomes.

## Methods

### Bacterial strains and pre-culturing procedures

Four strains were selected as inoculants for the soil interactome experiments: *P. veronii* 1YdBTEX2, a toluene, benzene, *m-* and *p-*xylene degrading bacterium isolated from contaminated soil^31^; *P. putida* F1, a benzene-, ethylbenzene- and toluene-degrading bacterium from a polluted creek^33^; *P. protegens* CHA0, a bacterium with plant-promoting character as a result of secondary metabolite production^34^; and *E. coli* MG1655^45^, as a typical non-soil dwelling bacterium. Variants of the four strains were deployed that constitutively express mCherry fluorescent protein.

*P. veronii* 1YdBTEX2 was tagged with a single-copy chromosomally inserted mini-Tn*7* transposon carrying a P_tac_–*mCherry* cassette (Pve, strain 3433) as described in Ref^46^. *P. putida* F1 was tagged with the same cassette but using mini-Tn*5* chromosomal delivery (Ppu, strain 5791). *P. protegens* CHA0 mini-Tn*7*:P_tac_-*mCherry* (Ppr, strain 6434) has been described previously^47^ and was kindly provided by Christoph Keel. mCherry expression in *E. coli* MG1655 was achieved from the same P_tac_-*mCherry* present on plasmid pME6012^48^ (Eco, strain 4514). All strains were kept at –80°C for long term storage and plated freshly for each experiment on nutrient agar (Oxoid Ltd.) containing the appropriate antibiotic for selection of the P_tac_-*mCherry* construct, from where individual colonies were transferred to liquid cultures.

Pve and Ppu colonies from nutrient agar were restreaked on 21C minimal media (MM)^49^ agar with toluene as sole carbon source provided through the vapour phase in a desiccator as described elsewhere^30^. A single colony of each after 48 h incubation at 30 °C was subsequently inoculated into 10 ml of liquid MM containing 5 mM sodium succinate as sole carbon source. Ppr and Eco colonies from selective nutrient agar plates were directly transferred to liquid MM with 5 mM succinate. Pve, Ppu and Ppr cultures were grown for 24 h at 30°C with rotary shaking at 180 rpm. Eco cultures were incubated at 37°C for 24 h with rotary shaking at 180 rpm. After 24 h, the cells were harvested and washed for bead encapsulation, as described below.

### Soil resident microbes

We chose sand as the source of the microbial community (which was hereafter named *soil community* or SC). The sand was collected fresh for each experiment from a beach of St. Sulpice near Lake Geneva (GPS coordinates: 46.508032 N, 6.544050 E) as described in Moreno et al^29^. Of note is that the sand was taken at different seasons and sampling times and may thus have carried slightly different starting communities and cell densities. The sand was sieved through 2 mm^2^ pores to remove large particles. The sieved sand was stored at room temperature and used within 7 days for extraction of resident microbial cells.

Microbial cells were extracted from four times 200 g of sand, each aliquot transferred in a 1 litre conical flask. Each 200 g of sand was submerged in 400 ml of 21C minimal media salts (MMS) (containing, per litre: 1 g NH_4_Cl, 3.49 g Na_2_HPO_4_·2H_2_O, 2.77 g KH_2_PO_4_, pH 6.8). Flasks were incubated at 25°C under rotary shaking at 120 rpm for one hour. The sand was allowed to settle and the supernatant was decanted into a set of 50 ml Falcon tubes, which were centrifuged at 800 rpm in an A-4-81 rotor in a 5810R centrifuge (Eppendorf AG.) for 10 min to precipitate heavy soil particles. Supernatants were decanted into clean 50 ml Falcon tubes and centrifuged at 4000 rpm for 30 minutes to pellet cells. The supernatants were carefully discarded, and the cell pellets were resuspended and pooled (i.e., from the initial 800 g of sand) in one tube using 5 ml of MMS. The pooled liquid suspension was further sieved through a 40–µm Falcon cell strainer (Corning Inc.) in order to remove any particles and large eukaryotic cells that may obstruct flow cytometry analysis (see below). A small proportion of the sieved liquid suspension was used for quantification of the extracted cell numbers (see below); the remainder was used within 12 h for bead encapsulation (see below). With this gentle method we extracted approximately 3 ×10^5^ cells g of sand^−1^.

### Flow cytometry cell counting

Cell numbers in suspensions of inoculants and extracted soil communities were counted by flow cytometry. Liquid cultures of the inoculant (10 ml) were centrifuged at 5000 rpm in a F-34-6-38 rotor in a 5804R centrifuge (Eppendorf AG) for 10 min at room temperature. The supernatant was discarded and the cell pellet was resuspended in 10 ml of MMS. The suspension was again centrifuged as before, supernatant was discarded and cells were resuspended in 10 ml of MMS. The inoculant suspension was then diluted 100 times in MMS and aspired at 66 µl min^−1^ on a Novocyte flow cytometer (ACEA Biosciences, USA). Inoculant cells were identified on the basis of their PE-Texas Red-H signal (channel voltage set at 592 V) representative for the mCherry fluorescence. An aliquot of the pooled SC cell suspension was diluted 100 times in MMS, and cells were stained by addition of SYTO-9 (1 µM final concentration), and incubation in the dark at room temperature for one hour. An aliquot of 30 µl was aspired on the flow cytometer and cells were counted above the FSC-H threshold of 500 and identified on the basis of their FITC-H signal (channel voltage at 441 V).

### Agarose bead encapsulation

Quantified inoculant and SC cells were mixed in 1:1 ratio in a 1 ml microcentrifuge tube, such that both contained approximately between 2×10^7^ and 10^8^ cells ml^−1^. As controls, batches of the inoculant-only or SC-only suspensions were used, each again between 2×10^7^ to 10^8^ cells ml^−1^.

To prepare beads in a size range of 40–70 µm, we used a procedure of rapid mixing of agarose– cell solution with pluronic acid in dimethylpolysiloxane and subsequent cooling and sieving. The whole procedure was carried out in a room maintained at 30°C and near a gas flame to maintain antiseptic conditions. 1% (*w*/*v*) low melting agarose (Eurobio ingen, France) was prepared in PBS solution (PBS contains per L H_2_O: 8 g NaCl, 0.2 g KCl, 1.44 g Na_2_HPO_4_, 0.24 g KH_2_PO_4_, pH 7.4) and dissolved by heating in a microwave. The molten agarose solution was cooled down, aliquoted to 1 ml batches in eppendorf vials and equilbrated in a 37°C water bath. Separately, 15 ml of dimethylpolysiloxane (Sigma-Aldrich) was poured in a 30 ml glass test tube. 1 ml of the 37°C– agarose solution was mixed with 30 µl of pluronic acid (Sigma-Aldrich) by vortexing at highest speed (Vortex-Genie 2, Scientific Industries, Inc.) for a minute. Into this mixture of agarose and pluronic acid, 200 µl of prepared cell suspension at 0.2–1.0 × 10^8^ cells ml^−1^ (inoculant only, SC only, or the mix of SC plus inoculant) was pipetted, and vortexed again at highest speed for another minute. 500 µl of this mixture was added drop-wise into the glass tube with dimethylpolysiloxane that was being vortexed at maximum speed. Vortexing was continued for two min. The tube was then immediately plunged into crushed ice and allowed to stand for a minimum of 10 min. After this, the total content of the tube was transferred into a 50 ml Falcon tube. The tube was centrifuged for 10 min at 2000 rpm using an A-4-81 swinging-bucket rotor (Eppendorf). The oil was carefully decanted, retaining the beads pellet. 15 ml of sterile PBS was added to the pellet, and the beads were resuspended by vortexing at a speed set to 5. The tubes were again centrifuged at 2000 rpm for 10 min, and any visible oil phase on the top was removed using a pipette. The process was repeated once more to remove any visible oil phase. Beads of diameter between 40–70 µm were then recovered by passing the PBS–resuspended bead content of the tube first over a 70–µm cell strainer (Corning Inc.). A further 5 ml of PBS was added to the cell strainer to flush remaining beads (<70–µm) into the filtrate. The collected bead filtrate was subsequently passed over a 40– µm cell strainer (Corning Inc.) to remove beads smaller than 40 µm. Recovered beads on the sieve were washed with an additional 5 ml of PBS, and any smaller beads in the filtrate that stuck to the bottom side of the cell strainer were gently removed by absorption with a Whatman 3M filter paper. After this, the sieve was inverted and placed on top of a clean 50 ml Falcon tube. 1.5 ml of incubation medium (MM with the respective carbon substrates, see below) was used to collect the beads from the sieve into the tube. Per interactome mixture, two tubes were prepared in parallel, which were pooled in the same final Falcon tube to yield a total volume of 3 ml that was split in three aliquots of 1 ml, to have triplicate interactome incubations. The encapsulation procedure produced ∼1.2×10^6^ beads per ml, with an effective volume of 10% of the total volume of the liquid phase in the incubations.

### Bead incubations

Four different carbon regimes were imposed as listed in Table 1. Toluene was used as example of an inoculant-selective substrate (Pve and Ppu), which we assumed would be poorly utilisable by the resident soil microbes and give selective benefit to the inoculant. Succinate, mixed carbon substrates (‘Mixed-C’), and sand extract (see below), were used as generally utilisable substrates for both inoculants and soil microbes. The external substrate concentration was limited to 0.1 mM (mixed-C) to avoid overgrowth of microcolonies inside the beads, which would lead to cell escape and their subsequent proliferation outside the beads. Experimental tests at higher succinate concentrations confirmed that overgrowth, cell escape and outside growth frequently occurred above 0.5 mM succinate.

Toluene was provided by partitioning from an oil phase. We diluted pure toluene 1000 times in 2,2’,4,4’,6,8,8’-heptamethylnonane (Sigma Aldrich) and added 0.2 ml of this solution to each vial with 1 ml bead suspension. A further 4 ml of MM was added to the vials.

Mixed-C solution was prepared by dissolving 16 individual compounds (Table S2) in milliQ-water (Siemens Labostar) in equimolar concentration such that the total carbon concentration of the solution reached 10 mM C. These compounds are also listed in EcoPlates™ (Biolog Hayward, CA, USA) and have been previously used as soil representative substrates ^25^. In the bead incubations, the mixed-C was diluted to 0.1 mM C final concentration in MM (total volume per vial again 5 ml). Sand extract was prepared by extraction with pre-warmed (70°C) sterile milliQ-water. A quantity of 100 g sand was mixed with 200 ml milliQ-water in a 250 ml Erlenmeyer flask and swirled on a rotatory platform for 15 min, after which it was subjected to 10 min sonication in an ultrasonic bath (Telesonic AG, Switzerland). Sand particles were sedimented and the supernatant was decanted, and passed through a 0.22–µm vacuum filter unit (Corning Inc.). This formed the ‘sand extract’, of which 4 ml was added directly to the 1 ml bead suspension in the vials.

For incubations with succinate, we added 4 ml of MM with 0.02 mM sodium succinate to each vial. Vials were incubated at 25 °C under rotary shaking at 110 rpm (to prevent too much settling of the beads), and were sampled for bacterial growth at regular time intervals (start, 6 h, 24 h, 48 h and 72 h).

### Bead sampling and microscopy

For sampling, the vials were removed from the incubator and beads were spun down at 1200 rpm using a swinging-bucket A-4-81 rotor (Eppendorf). An aliquot of 10 µl of bead suspension was carefully sampled from the bottom of the vials, mixed with 0.6 µl of 50 µM SYTO-9 solution to stain all cells, and incubated for 20 min at room temperature in the dark. Vials were placed back into the incubator. 5 µl sterile milli-Q water was added to the stained beads, and the complete aliquot (15 µl) was spread on a regular microscope glass slide to minimise aggregation of beads. A coverslip (24 x 50 mm) was placed gently avoiding air bubbles and excessive squeezing of the beads. Ten random positions on the slide were imaged with the 20× objective (NA 0.35) using an inverted AF6000 LX epifluorescence microscope system (Leica AG, Germany), equipped with a DFC350FXR2 camera. Every position was imaged in four sequential channels (phase contrast, 25 ms; mCherry, Y3–cube, 750 ms; SYTO-9, GFP–cube, 50 and 340 ms). The 50 ms–SYTO-9 channel exposure was used for analysis; the 340–ms exposure was used for verification of weak signals, if necessary. Images were recorded as 16-bit TIF-files and further processed using a custom MatLab routine (described below).

### Microscopy image analysis

A custom Matlab image processing and analysis routine was developed to segment beads and microcolonies inside beads from the image-series (Fig. S1), to identify and differentiate inoculant cell colonies (visible in mCherry and SYTO9) and colonies originating from sand cells (only visible in SYTO9). The mean fluorescence intensity and area of each identified microcolony type was quantified, from which the number of microcolonies within beads and their geometric distances were calculated.

For each time-point and experimental replicate, the phase contrast, mCherry, and SYTO-9 images were read using the *imread* function built in MatLab (version 2016b, MathWorks inc., USA). To identify the beads on each image, sharp changes in intensity were detected in the phase-contrast images using the *edge* function. Individual beads within a specific radius range were then identified using the *imfindcircles* function. In the next step, the microcolonies inside each bead were identified, by thresholding and segmenting the mCherry and SYTO-9 images, exclusively within the identified bead areas. mCherry and SYTO-9 images were further aligned to identify microcolonies in SYTO-9 having mCherry signal, which corresponds to the inoculant. Overlapping signals were considered to originate from an inoculant colony if the area overlap between two channels was greater than 30%. Else, the areas were considered to consist of both inoculant and SC cells. All microcolonies were thus differentiated as corresponding to inoculant (mCherry plus SYTO-9 signal) or SC (SYTO-9 only), after which their area, fluorescence intensity and inter-particle distance (within the bead) were calculated.

Results were summarized for each incubation and time point to comprise the following information: (i) total number of beads for each of the treatments (SC–cell only, inoculant only, or SC plus inoculant); (ii) the product of the particle pixel area times its SYTO-9 fluorescence intensity (we refer to this as productivity); (iii) the number of particles per bead and bead types (i.e., being only SC, only inoculant, or both); (iv) the summed averaged per-bead productivity of the different experimental treatments (i.e., type of inoculant, type of carbon substrate, time effect); and (v) the individual productivities of beads with pairs of SC-inoculant, or SC-SC. The MatLab routines are provided as Supplementary Code.

### Statistical testing

Aggregate productivities in incubations and per category (inoculant, SC or both) or substrate were compared in ANOVA, with significance testing inferred post hoc according to Tukey. Maximum aggregate productivities across multiple different substrates (yielding different biomass) were compared in a non-parametric rank sum Wilcoxon test. Given their non-normal nature, bead productivity distributions were globally compared non-parametrically with the Fisher test. Further parametric (t-test and ANOVA) testing on bead productivity distributions was conducted using boosted averages (>75^th^ percentile) and >95^th^ percentile sums.

## ACKNOWLEDGMENTS

This work was supported by SystemsX.ch, and evaluated by the Swiss National Science Foundation, within grant 2013/158 (Design and Systems Biology of Functional Microbial Landscapes ‘MicroScapesX’).

## SUPPLEMENTARY MATERIAL

**Supplementary Fig. S1 Microcolony development in beads.** A. Starting cells in agarose beads of Pve, Eco, Ppr, Ppu and sand community (SC), stained by SYTO-9. B. Microcolonies of Pve, Eco, Ppr and Ppu after 24-72 h (constitutive mcherry signal) and of SC (SYTO-9).

**Supplementary Fig. S2 Maximum productivities by *P. veronii* and SC across multiple independent incubation series.** Squares represent maximum productivity at any of the (triplicate) incubations at time points (24, 48 and 72 h) of Pve (black) or SC (light blue) separately, or in combination (green). MIXC1–C5, five independent repetitions of incubations with mixed C substrates; SUCC1–2, two independent repetitions with 0.1 mM succinate; TOL1–4, four repetitions with toluene (supplied through the vapor phase). Note that SC–cells were extracted from freshly taken sand material at different occasions (seasons) and thus may have different starting cell compositions. See main text for statistical tests.

**Supplementary Fig. S3 Productivity shifts in inoculant–SC pairs compared to either inoculant or SC alone.** Productivity is defined as the product of particle area and fluorescence intensity. Distributions per category plotted using the gaussian density kernel on log scale. Substrates and inoculants abbreviated as before.

**Supplementary Fig. S4 Summed productivities among inoculant–SC pairs of which the SC productivity was above the 95^th^ percentile of the SC productivity separately after 24 h on the same carbon regime.** Distributions show log-scale productivities of the exclusive inoculant (Ino)–SC pairs (in green), compared to all inoculant–SC pairs (magenta) or inoculant alone (light brown). Compare to Figure 5 and Fig. S5. mixC, mixed carbon substrates; TOL, toluene. Inoculant abbreviations as before. P-values in Fisher’s exact test of comparing normalized histogram distributions.

**Supplementary Fig. S5 Productivities of inoculant–SC pairs.** A. Pair-wise productivity on mixed carbon substrates of co-occurring *P. protegens* (Ppr) and sand community (SC) within the same bead (colored bubbles), compared to productivity of Ppr or SC in separate individual incubations. B. As A, but for *E. coli* (Eco). C. as A, but for *P. putida* (Ppu) and toluene. D. as A, but for *P. veronii* and sand extract. Productivities are displayed on log-axes. Green lines indicate the 95^th^ percentile productivity of the individual incubation at t=24 h. Bubble diameters represent the Euclidian distance from the origin and are a relative measure of the microcolony sizes. Percentages indicate the proportion of beads of the total, falling above the respective 95^th^ percentile threshold. Values reported in Table 2.

## Supplementary Code

MatLab subroutines for microcolony in bead analysis.

