## Supplemental Figures 1-5, Supplemental code for "Close-Range Interactions Favor Growth in Random-Paired Extracted Soil Bacteria"

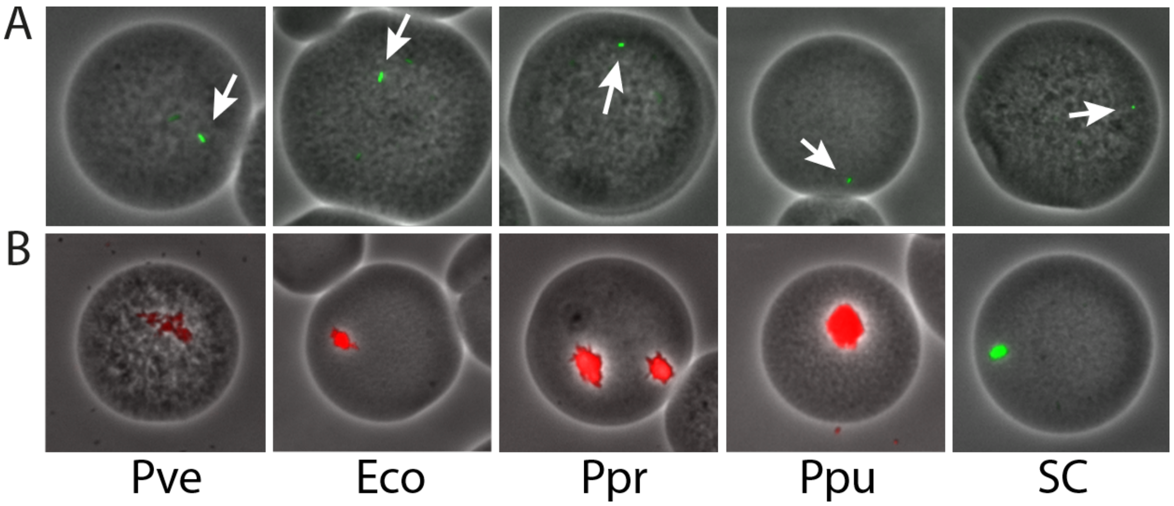


**Supplementary Fig. S1 Microcolony development in beads.** A. Starting cells in agarose beads of Pve, Eco, Ppr, Ppu and sand community (SC), stained by SYTO-9. B. Microcolonies of Pve, Eco, Ppr and Ppu after 24-72 h (constitutive mcherry signal) and of SC (SYTO-9).

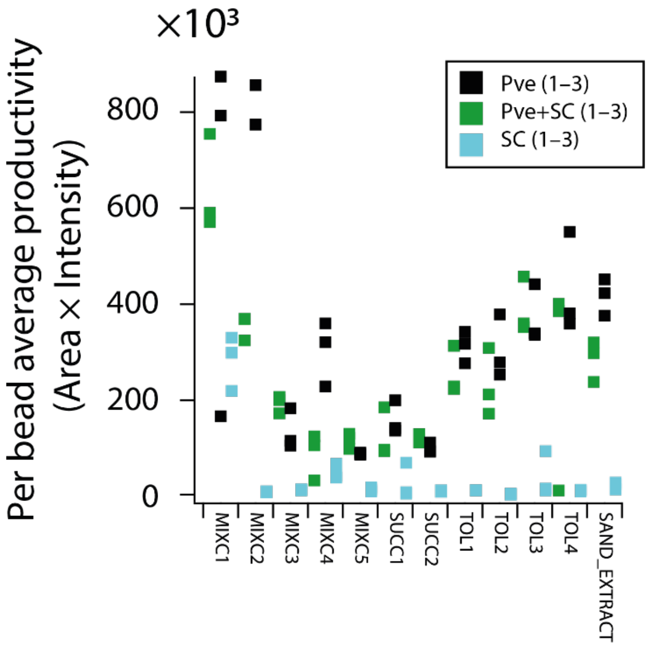


**Supplementary Fig. S2 Maximum productivities by *P. veronii* and SC across multiple independent incubation series.** Squares represent maximum productivity at any of the (triplicate) incubations at time points (24, 48 and 72 h) of Pve (black) or SC (light blue) separately, or in combination (green). MIXC1–C5, five independent repetitions of incubations with mixed C substrates; SUCC1–2, two independent repetitions with 0.1 mM succinate; TOL1–4, four repetitions with toluene (supplied through the vapor phase). Note that SC–cells were extracted from freshly taken sand material at different occasions (seasons) and thus may have different starting cell compositions. See main text for statistical tests.

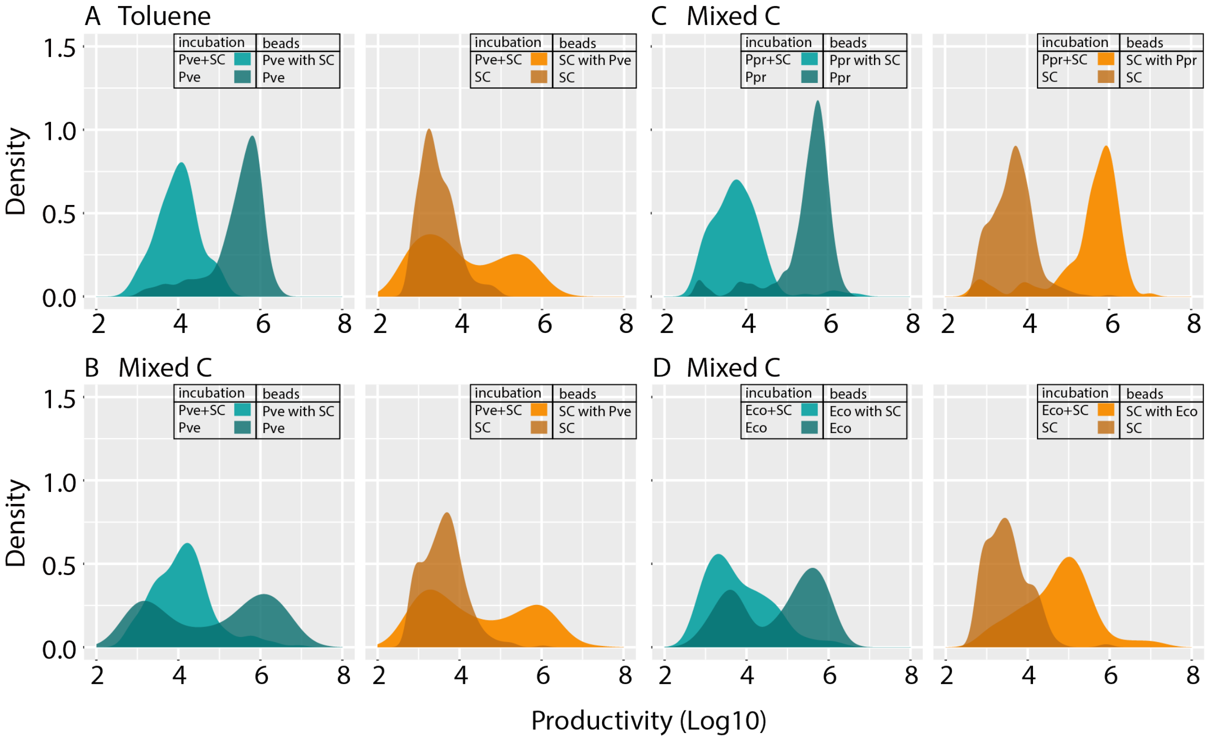


**Supplementary Fig. S3 Productivity shifts in inoculant–SC pairs compared to either inoculant or SC alone.** Productivity is defined as the product of particle area and fluorescence intensity. Distributions per category plotted using the gaussian density kernel on log scale. Substrates and inoculants abbreviated as before.

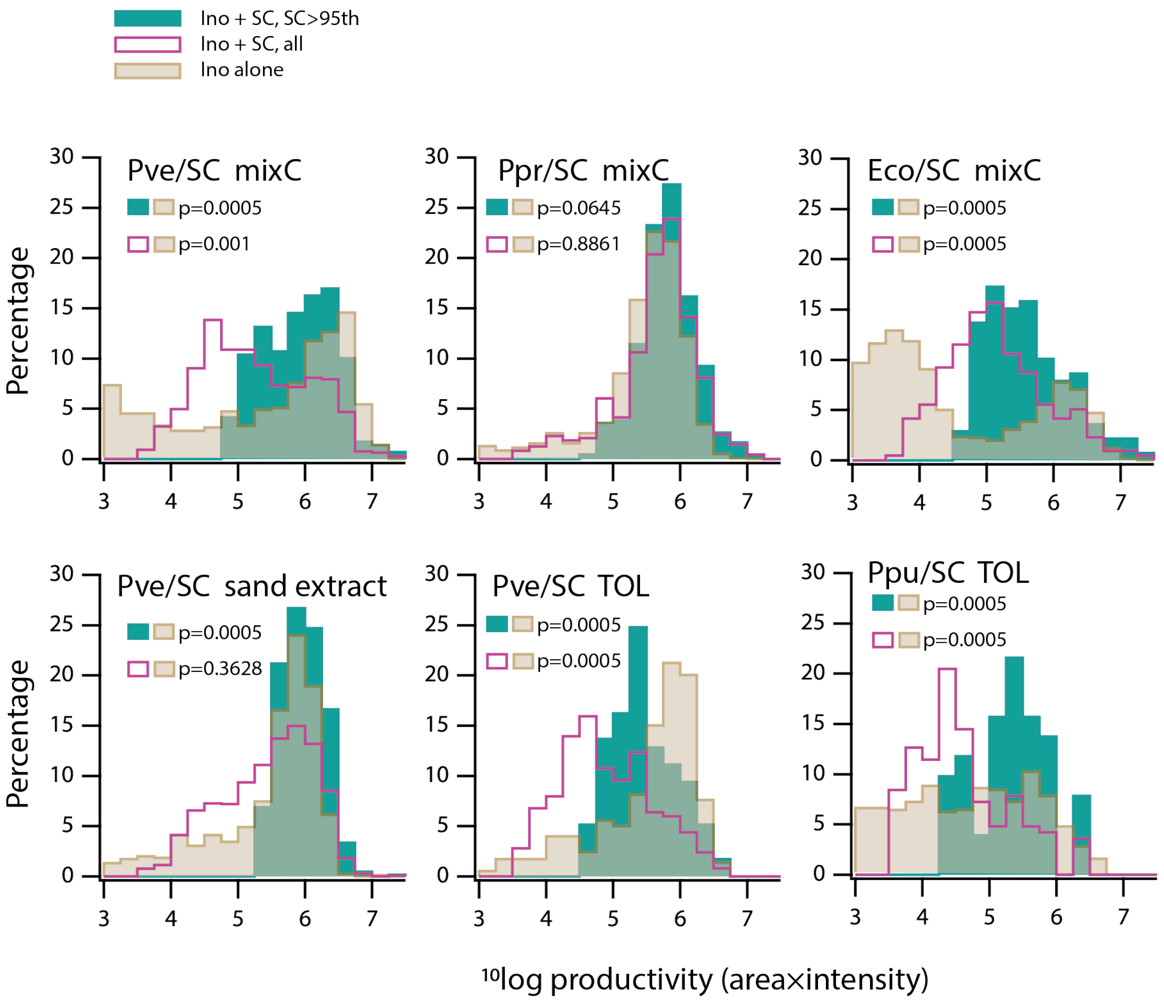


**Supplementary Fig. S4 Summed productivities among inoculant–SC pairs of which the SC productivity was above the 95$^{th}$ percentile of the SC productivity separately after 24 h on the same carbon regime.** Distributions show log-scale productivities of the exclusive inoculant (Ino)–SC pairs (in green), compared to all inoculant–SC pairs (magenta) or inoculant alone (light brown). Compare to Figure 5 and Fig. S5. mixC, mixed carbon substrates; TOL, toluene. Inoculant abbreviations as before. P-values in Fisher's exact test of comparing normalized histogram distributions.

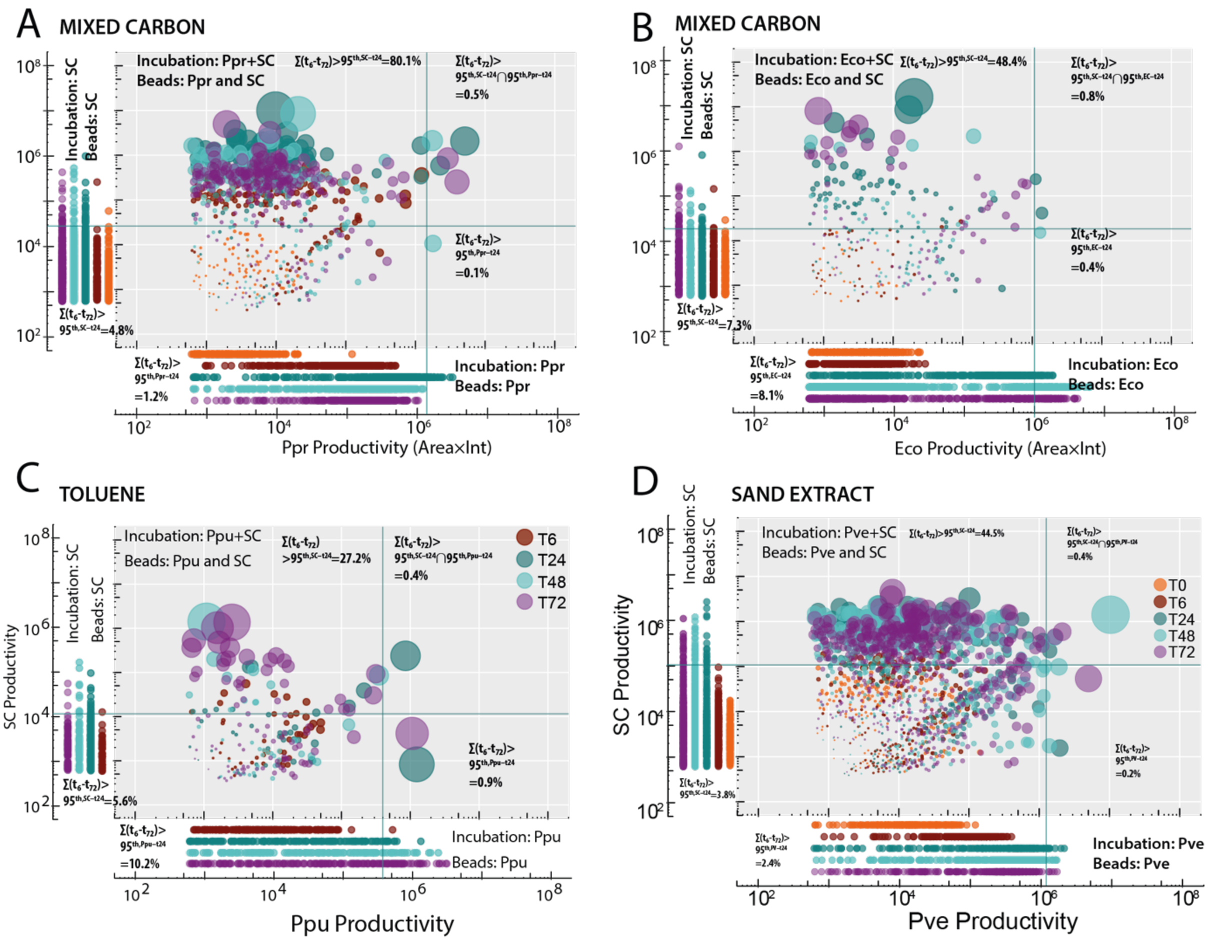


**Supplementary Fig. S5 Productivities of inoculant–SC pairs.** A. Pair-wise productivity on mixed carbon substrates of co-occurring *P. protegens* (Ppr) and sand community (SC) within the same bead (colored bubbles), compared to productivity of Ppr or SC in separate individual incubations. B. As A, but for *E. coli* (Eco). C. as A, but for *P. putida* (Ppu) and toluene. D. as A, but for *P. veronii* and sand extract. Productivities are displayed on log-axes. Green lines indicate the 95$^{th}$ percentile productivity of the individual incubation at t=24 h. Bubble diameters represent the Euclidian distance from the origin and are a relative measure of the microcolony sizes. Percentages indicate the proportion of beads of the total, falling above the respective 95$^{th}$ percentile threshold. Values reported in Table 2.

Supplementary Code 1.

Authors: S. Pelet, N. Hadadi, M. Dubey and J. R. van der Meer
Department of Fundamental Microbiology, University of Lausanne
1015 Lausanne, Switzerland

This file contains four subfiles that analyse the bead images with microcolonies, based on two colors (GFP, mCherry) and a phase contrast image.
Individual parts can be run in MatLab.
Part 1 contains the code to define the beads, define the microcolonies and measure the fluorescence area and intensity of the microcolonies within the beads. It also separates the 'inoculant' (based on its mCherry signal) from the 'sand cells' (which only carry the Syto9 stain). Part 1 writes a table for every set of image folders with a single experiment.

Part 2-4 collects all inputs, gather all columns relevant for sand cell interactions and write a summary output readable by Excel.

```
%%PART 1

function Table = SegmentBeads_6(Folder)
close all
%% Analyses the entire experiment. Generates files as described below in each replicate folder containing images.
%%NH\June 2017: Load images from main folder\time points\experiments (run the routine for all the experiments simultaniously, instead of one by one)
Main='/folderX/folderY/ExperimentFolder/'
Timepoints=dir([Main,'T*']);

%inside each time point
for t=1:length(Timepoints)
   Timepoint=fullfile(Main, Timepoints(t).name);
Experiment=dir(Timepoint);
Experiment = Experiment(~strncmpi('.', {Experiment.name}, 1));

%inside each experiment
for e=1:length(Experiment)
   Folder=fullfile(Timepoint, Experiment(e).name,'/');
PhaseImgList = dir([Folder,'*_ch00.tif']);
IterGFP = 0;
IterCherry = 0;
%%inside each image
for F = 1:length(PhaseImgList)
   PhaseImg = double(imread(fullfile(Folder, PhaseImgList(F).name)));
   SYTOname = [PhaseImgList(F).name(1:end-5), '2.tif'];
   SYTOImg = double(imread(fullfile(Folder, SYTOname)));
   Cherryname = [PhaseImgList(F).name(1:end-5), '1.tif']
   CherryImg = double(imread(fullfile(Folder, Cherryname)));

%% Parameters

BeadsRadii = [30, 100];          %Min and max size of bead radius
Sensitivity = 0.93;              %Default 0.85

SYTOthresh = 300;                %Intensity threshold for SYTO image %Set to zero for automated threshold
Cherrythresh = 220;              %Intensity threshold for Cherry image
```

```
FillObjects = 1;                %Set to one if objects need to be filled in after edge detection

threshold_For_SC=0.3;           %added by NH for analyzing data from excel file

areathresh = 2;                 %added by Manu and NH for removing the very small pixels

%% Segment Phase image

%detect edges
[PC_edge, Thresh] = edge(PhaseImg, 'Canny');
[PC_edge, Thresh] = edge(PhaseImg, 'Canny', Thresh*2);

if FillObjects == 1

%Connect pixels in edges
PC_bridge = bwmorph(PC_edge, 'bridge');
%Close image
SE = strel('disk', 5);
PC_bridge = imclose(PC_bridge, SE);
%Fill image
PC_fill = imfill(PC_bridge, 'holes');

PC_segment = PC_fill;
else
   PC_segment = PC_edge;
end
% figure(2)
% imagesc(PC_segment)

%Hough transform to indentify circles in image
[Centers, Radii] = imfindcircles(PC_segment,BeadsRadii, 'ObjectPolarity', 'bright',...
   'Sensitivity', Sensitivity);

%Display circles on Phase contrast image
% figure(1)
% imagesc(PhaseImg)
% hold on
% viscircles(Centers, Radii,'EdgeColor','b');

%Sort Radius to start with larger cells
[Radii, RadOrder] = sort(Radii, 'descend');
Centers = Centers(RadOrder,:);
NbBeads = length(Radii);

%Initialize variables
BeadImg = zeros(size(PhaseImg));
IterCell = 0;
%Loop through all beads
for B = 1:NbBeads

   % find pixels within circle given by radius and center
   %Get coordinates of 100 perimeter pixels based on center and radius
   Angle = [0 : (2 * pi / 100) : (2 * pi)];
```

```
    XPerim = Radii(B) * cos(Angle) + Centers(B,1);
    YPerim = Radii(B) * sin(Angle) + Centers(B,2);

    %Create matrix containing all coordinates in the square containing
    %the circle.
    [XGr,YGr] = meshgrid([floor(Centers(B,1)-1.1*Radii(B)) : ceil(Centers(B,1)+1.1*Radii(B))],...
        [floor(Centers(B,2)-1.1*Radii(B)) : ceil(Centers(B,2)+1.1*Radii(B))]);
    %Make sure XGr and YGr are not beyond the limit of the image
    XGr(XGr<=0) = 1;
    YGr(YGr<=0) = 1;
    YGr(YGr>size(PhaseImg,1)) = size(PhaseImg,1);
    XGr(XGr>size(PhaseImg,2)) = size(PhaseImg,2);
    %Find pixels contained in the perimeter
    InBead = inpolygon(XGr(:),YGr(:),XPerim,YPerim);
    %Convert X-Y subscripts to indices
    InInd = sub2ind(size(PhaseImg),YGr(InBead),XGr(InBead));
    %Test if center lies whitin a previously identified bead
    if BeadImg(round(Centers(B,2)), round(Centers(B,1))) ~= 0
        %If center is within previous identified circle: Merge with this
        %circle
        BeadImg(InInd) = BeadImg(round(Centers(B,2)), round(Centers(B,1)));
    else
        %Other wise create a new bead object
        IterCell = IterCell+1;
        BeadImg(InInd) = IterCell;
    end
end

% figure(4)
% imagesc(BeadImg)

%% Segment SYTO image
%Set pixels outside of segmented beads to the median value of these pixels
% => this removes SYTO signals outside of beads
SYTOImg(BeadImg==0) = median(SYTOImg(BeadImg==0));


%Normalize the image
NormSYTO = (SYTOImg-min(SYTOImg(:)))./(max(SYTOImg(:))-min(SYTOImg(:)));
NormSYTO = imadjust(NormSYTO, [0.01, 0.90], []);
if SYTOthresh == 0
    %Calculate threshold for segmentation
    Tresh = graythresh(NormSYTO);
    %Generate BW image where object pixels are set to 1
    BWSyto = im2bw(NormSYTO,Tresh);
else
    BWSyto = zeros(size(SYTOImg));
    BWSyto(SYTOImg>SYTOthresh) = 1;
end

%Calculate properties of all objects
CC = bwconncomp(BWSyto);
SYTOStats = regionprops(CC, SYTOImg, 'Area','PixelIdxList', 'MeanIntensity', 'Centroid');
% figure(15)
```

```
% imagesc(BWSyto); title('Syto segmentation')

%% Segment mCherry image
%Set pixels outside of segmented beads to the median value of these pixels
% => this removes mCherry signals outside of beads
CherryImg(BeadImg==0) = median(CherryImg(BeadImg==0));
%Normalize the image
NormCherry = (CherryImg-min(CherryImg(:)))./(max(CherryImg(:))-min(CherryImg(:)));
NormCherry = imadjust(NormCherry, [0.01, 0.90], []);

if Cherrythresh == 0
   %Calculate threshold for segmentation
   Tresh = graythresh(NormCherry);
   %Generate BW image where object pixels are set to 1
   BWCherry = im2bw(NormCherry,Tresh);
else
   BWCherry = zeros(size(CherryImg));
   BWCherry(CherryImg>Cherrythresh) = 1;
end

%Calcualte properties of all objects
CC = bwconncomp(BWCherry);
CherryStats = regionprops(CC, CherryImg, 'Area','PixelIdxList', 'MeanIntensity', 'Centroid');

% figure(16)
% imagesc(BWCherry) ; title('Cherry segmentation')

%% Compare Cherry and SYTO segmentations
%Run through all SYTO objects

for S = 1:length(SYTOStats)
   %Find the indices of objects in the bead image
   BeadNum = unique(BeadImg(SYTOStats(S).PixelIdxList));
   %Check if object belongs to a single bead
   if length(BeadNum) == 1
      %Increase iteration value
      IterGFP = IterGFP+1;
      %Save object area
      SytoArea(IterGFP,1)  = SYTOStats(S).Area;
      %Save object mean intensity
      SytoIntensity(IterGFP,1)  = SYTOStats(S).MeanIntensity;
      SytoCentroidX(IterGFP,1)  = SYTOStats(S).Centroid(1);
      SytoCentroidY(IterGFP,1)  = SYTOStats(S).Centroid(2);
      %Save object Bead Number
      BeadNumber(IterGFP,1) = BeadNum;

      %Calculate overlap with Cherry pixels
      %Find pixels belonging to SYTO object which are Red
      OverlapPixels = find(BWCherry(SYTOStats(S).PixelIdxList)==1);
      CherryOverlapArea(IterGFP,1) = length(OverlapPixels);
      CherryOverlapIntensity(IterGFP,1) = mean(CherryImg(OverlapPixels));
      %Calculate extent of colocalization
      CherryColoc(IterGFP,1) = CherryOverlapArea(IterGFP)/SytoArea(IterGFP);
      SandCellArea(IterGFP,1) = SytoArea(IterGFP) - CherryOverlapArea(IterGFP);
```

```
        ImageNameGFP{IterGFP,1} = PhaseImgList(F).name;
    end
end

%% Compare Cherry segmentations
%Run through all Cherry objects

for S = 1:length(CherryStats)
    %Find the indices of objects in the bead image
    BeadNum2 = unique(BeadImg(CherryStats(S).PixelIdxList));
    %Check if object belongs to a single bead
    if length(BeadNum2) == 1
        %Increase iteration value
        IterCherry = IterCherry+1;
        %Save object area
        CherryArea(IterCherry,1)  = CherryStats(S).Area;
        %Save object mean intensity
        CherryIntensity(IterCherry,1)  = CherryStats(S).MeanIntensity;
        CherryCentroidX(IterCherry,1)  = CherryStats(S).Centroid(1);
        CherryCentroidY(IterCherry,1)  = CherryStats(S).Centroid(2);
        %Save object Bead Number
        BeadNumber2(IterCherry,1) = BeadNum2;
        ImageNameCherry{IterCherry,1} = PhaseImgList(F).name;
    end
end

%% Img Display

% figure(F*100)
% RGBsegment(:,:,1) = BWCherry;
% RGBsegment(:,:,2) = BWSyto;
% BWBeads = zeros(size(BeadImg));
% BWBeads(BeadImg>0) = 0.5;
% RGBsegment(:,:,3) = BeadImg;
% image(RGBsegment)
%
% figure(F*100+1)
%
% RGBImg(:,:,1) = NormCherry;
%
% RGBImg(:,:,2) = NormSYTO;
%
% NormPhase = (PhaseImg-min(PhaseImg(:)))./(max(PhaseImg(:))-min(PhaseImg(:)));
% NormPhase = imadjust(NormPhase);
% RGBImg(:,:,3) = NormPhase.*0.5;
% image(RGBImg)
% drawnow
end

%% Save
%Create table
TableGFP = table(ImageNameGFP, BeadNumber, SytoCentroidX, SytoCentroidY, SytoArea, SytoIntensity,
SandCellArea, CherryOverlapArea, CherryOverlapIntensity, CherryColoc);
```

```
%Create table
TableCherry = table(ImageNameCherry, BeadNumber2, CherryArea, CherryIntensity,
CherryCentroidX,CherryCentroidY);
%Save in the folder

%remove lines from the table that the area is below the defined
%threshold (Manu and NH)
toDelete = TableGFP .SytoArea < areathresh;
TableGFP(toDelete,:) = [];

TableGFP_1=sortrows(TableGFP,2,'ascend');
TableGFP_2=sortrows(TableGFP_1,1,'ascend');

%% analyzing data and creating a summary table %NH/May2017

BeadNumber=TableGFP_2.BeadNumber;
CherryColoc=TableGFP_2.CherryColoc;
CherryOverlapArea=TableGFP_2.CherryOverlapArea;
SandCellArea=TableGFP_2.SandCellArea;
SytoArea=TableGFP_2.SytoArea;
SytoIntensity=TableGFP_2.SytoIntensity;

[a,b]=size(BeadNumber);

Count_NrBeads_inImage=zeros(a,b);
Count_Total_NrBeads=zeros(a+1,b);
ParticleNr=zeros(a,b);
BeadSumCheOver=zeros(a,b);
BeadSumSytoArea_times_Int=zeros(a,b);
beadsWithSConly=zeros(a,b);
list_AtimesI_SC_only=zeros(a,b);
BeadSumSandArea=zeros(a,b);
list_for_counting_PV=zeros(a,b);
list_for_Area_times_Int_PV=zeros(a,b);
SC_A_times_I_inbeads_with_PV=zeros(a,b);
list_SC_A_times_I_per_bead=zeros(a,b);
Count_Beads_with_SC_and_PV=zeros(a,b);
list_SC_A_times_I_inbeads_with_PV=zeros(a,b);
list_PV_A_times_I_inbeads_with_SC=zeros(a,b);
PV_A_times_I_inbeads_with_SC=zeros(a,b);
list_for_Area_times_Int_PV_with_SC=zeros(a,b);

Count_NrBeads_inImage(1,1)=1;
Count_Total_NrBeads(1,1)=1;
ParticleNr(1,1)=1;
beadsWithSConly(1,1)=0;
list_for_counting_PV(1,1)=0;
Count_Beads_with_SC_and_PV(1,1)=0;

for i=1:a-1

    if  BeadNumber(i+1,1)> BeadNumber(i,1)
        Count_NrBeads_inImage(i+1,1)=Count_NrBeads_inImage(i,1)+1;
    elseif BeadNumber(i+1,1)< BeadNumber(i,1)
```

```
        Count_NrBeads_inImage(i+1,1)=1;
    else
        Count_NrBeads_inImage(i+1,1)=Count_NrBeads_inImage(i,1);
    end

    if  Count_NrBeads_inImage(i+1,1)== Count_NrBeads_inImage(i,1)
        Count_Total_NrBeads(i+1,1)=Count_Total_NrBeads(i,1);
    else
        Count_Total_NrBeads(i+1,1)=Count_Total_NrBeads(i,1)+1;
    end

    if  Count_Total_NrBeads(i+1,1)== Count_Total_NrBeads(i,1)
        ParticleNr(i+1,1)=ParticleNr(i,1)+1;
    else
        ParticleNr(i+1,1)=1;
    end

    BeadSumCheOver(1,1)=CherryOverlapArea(1,1);

    if ParticleNr(i+1,1)> ParticleNr(i,1)
        BeadSumCheOver(i+1,1)=CherryOverlapArea(i+1,1)+BeadSumCheOver(i,1);
    else
        BeadSumCheOver(i+1,1)= CherryOverlapArea(i+1,1);
    end

    BeadSumSytoArea_times_Int(1,1)=SytoArea(1,1)*SytoIntensity(1,1);
    if ParticleNr(i+1,1)> ParticleNr(i,1)
        BeadSumSytoArea_times_Int(i+1,1)=BeadSumSytoArea_times_Int(i,1)+(SytoArea(i+1,1)*SytoIntensity(i+1,1));
    else
        BeadSumSytoArea_times_Int(i+1,1)=SytoArea(i+1,1)*SytoIntensity(i+1,1);
    end

end

Count_Total_NrBeads(end,1)=Count_Total_NrBeads(end-1,1);

for i=1:a-1
    if Count_Total_NrBeads(i+1,1)== Count_Total_NrBeads(i+2,1)
        beadsWithSConly(i+1,1)=-1;
    elseif Count_Total_NrBeads(i+1,1)< Count_Total_NrBeads(i+2,1)&& BeadSumCheOver(i+1,1)~=0
        beadsWithSConly(i+1,1)=0;
    elseif Count_Total_NrBeads(i+1,1)< Count_Total_NrBeads(i+2,1) && BeadSumCheOver(i+1,1)==0 &&
BeadSumCheOver(i+1,1)<=BeadSumCheOver(i,1)
        beadsWithSConly(i+1,1)=1;
    elseif Count_Total_NrBeads(i+1,1)< Count_Total_NrBeads(i+2,1) && BeadSumCheOver(i+1,1)==0 &&
BeadSumCheOver(i+1,1)> BeadSumCheOver(i,1)
        beadsWithSConly(i+1,1)=0;
    end
end

for i=1:a-1
    if beadsWithSConly(i)==1
        list_AtimesI_SC_only(i)=BeadSumSytoArea_times_Int(i);
    else
```

```
        list_AtimesI_SC_only(i)=0;
    end
    BeadSumSandArea(1,1)=SandCellArea(1,1);
    if ParticleNr(i+1,1)> ParticleNr(i,1) && CherryColoc(i+1)> threshold_For_SC
        BeadSumSandArea(i+1,1)=BeadSumSandArea(i,1);
    elseif ParticleNr(i+1,1)> ParticleNr(i,1) && CherryColoc(i+1) <= threshold_For_SC
        BeadSumSandArea(i+1,1)=BeadSumSandArea(i,1)+SandCellArea(i+1,1);
    elseif ParticleNr(i+1,1)<= ParticleNr(i,1) && CherryColoc(i+1)> threshold_For_SC
        BeadSumSandArea(i+1,1)=0;
    elseif ParticleNr(i+1,1)<= ParticleNr(i,1) && CherryColoc(i+1)<= threshold_For_SC
        BeadSumSandArea(i+1,1)=SandCellArea(i+1,1);
    end
end

for i=1:a-1
    if Count_Total_NrBeads(i+1,1)< Count_Total_NrBeads(i+2,1) && BeadSumSandArea(i+1,1)==0
        list_for_counting_PV(i+1,1)=1;
    elseif Count_Total_NrBeads(i+1,1)< Count_Total_NrBeads(i+2,1) && BeadSumSandArea(i+1,1)~=0
        list_for_counting_PV(i+1,1)=-1;
    else list_for_counting_PV(i+1,1)=0;
    end
end

for i=1:a-1
    if list_for_counting_PV(i,1)==1
        list_for_Area_times_Int_PV(i,1)=BeadSumSytoArea_times_Int(i,1);
    else
        list_for_Area_times_Int_PV(i,1)=0;
    end

    if CherryColoc(i)< threshold_For_SC
        SC_A_times_I_inbeads_with_PV(i,1)=SytoArea(i,1)*SytoIntensity(i,1);
    else
        SC_A_times_I_inbeads_with_PV(i,1)=0;
    end
end

list_SC_A_times_I_per_bead(1,1)=SC_A_times_I_inbeads_with_PV(1,1);

for   i=1:a-1
    if ParticleNr(i+1,1)> ParticleNr(i,1)
        list_SC_A_times_I_per_bead(i+1,1)=list_SC_A_times_I_per_bead(i,1)+ SC_A_times_I_inbeads_with_PV(i+1,1);
    else
        list_SC_A_times_I_per_bead(i+1,1)=SC_A_times_I_inbeads_with_PV(i+1,1);
    end

    if beadsWithSConly(i+1,1)==0 && list_for_counting_PV(i+1,1)==1
        Count_Beads_with_SC_and_PV(i+1,1)=0;
    elseif beadsWithSConly(i+1,1)==0 && list_for_counting_PV(i+1,1)~=1
        Count_Beads_with_SC_and_PV(i+1,1)=1;
    else
        Count_Beads_with_SC_and_PV(i+1,1)=0;
    end
```

```
    if Count_Beads_with_SC_and_PV(i,1)==1
        list_SC_A_times_I_inbeads_with_PV(i,1)=list_SC_A_times_I_per_bead(i,1);
    else list_SC_A_times_I_inbeads_with_PV(i,1)=0;
    end

    if CherryColoc(i)>= threshold_For_SC
        list_PV_A_times_I_inbeads_with_SC(i,1)=SytoArea(i,1)*SytoIntensity(i,1);
    else
        list_PV_A_times_I_inbeads_with_SC(i,1)=CherryOverlapArea(i,1)*SytoIntensity(i,1);
    end
end

PV_A_times_I_inbeads_with_SC(1,1)=list_PV_A_times_I_inbeads_with_SC(1,1);

for   i=1:a-1

    if ParticleNr(i+1,1)> ParticleNr(i,1)

PV_A_times_I_inbeads_with_SC(i+1,1)=PV_A_times_I_inbeads_with_SC(i,1)+list_PV_A_times_I_inbeads_with_SC(
i+1,1);
    else PV_A_times_I_inbeads_with_SC(i+1,1)=list_PV_A_times_I_inbeads_with_SC(i+1,1);
    end

    if Count_Beads_with_SC_and_PV(i,1)==1
        list_for_Area_times_Int_PV_with_SC(i,1)= PV_A_times_I_inbeads_with_SC(i,1);
    else   list_for_Area_times_Int_PV_with_SC(i,1)=0;
    end
end

Total_nr_beads=max(Count_Total_NrBeads);
Total_nr_beads_with_SC_only=numel(beadsWithSConly(beadsWithSConly==1));
Total_nr_beads_with_PV_only=numel(list_for_counting_PV(list_for_counting_PV==1));
Total_nr_beads_with_SC_and_PV=numel(Count_Beads_with_SC_and_PV(Count_Beads_with_SC_and_PV==1));

SUM_A_times_I_SC_only=sum(list_AtimesI_SC_only);
SUM_A_times_I_PV_only=sum(list_for_Area_times_Int_PV);
SUM_A_times_I_SC_with_PV=sum(list_SC_A_times_I_inbeads_with_PV);
SUM_A_times_I_PV_with_SC=sum(list_for_Area_times_Int_PV_with_SC);

proportion_SC_beads=Total_nr_beads_with_SC_only/Total_nr_beads;
proportion_PV_beads=Total_nr_beads_with_PV_only/Total_nr_beads;
proportion_SC_and_PV_beads=Total_nr_beads_with_SC_and_PV/Total_nr_beads;

A_times_I_per_bead_forSC_alone=SUM_A_times_I_SC_only/Total_nr_beads_with_SC_only;
A_times_I_per_bead_forSC_inPV=SUM_A_times_I_SC_with_PV/Total_nr_beads_with_SC_and_PV;
A_times_I_per_bead_forPV_alone=SUM_A_times_I_PV_only/Total_nr_beads_with_PV_only;
A_times_I_per_bead_forPV_inSC=SUM_A_times_I_PV_with_SC/Total_nr_beads_with_SC_and_PV;

values=[Total_nr_beads, Total_nr_beads_with_SC_only,Total_nr_beads_with_PV_only,
Total_nr_beads_with_SC_and_PV,SUM_A_times_I_SC_only, SUM_A_times_I_PV_only,
SUM_A_times_I_SC_with_PV, SUM_A_times_I_PV_with_SC, proportion_SC_beads, proportion_PV_beads,
proportion_SC_and_PV_beads, A_times_I_per_bead_forSC_alone, A_times_I_per_bead_forSC_inPV,
A_times_I_per_bead_forPV_alone, A_times_I_per_bead_forPV_inSC];
values=values';
```

```
values=num2cell(values);
headers=
{'Total_nr_beads','Total_nr_beads_with_SC_only','Total_nr_beads_with_PV_only','Total_nr_beads_with_SC_and_PV',
'SUM_A_times_I_SC_only','SUM_A_times_I_PV_only','SUM_A_times_I_SC_with_PV','SUM_A_times_I_PV_with_
SC','proportion_SC_beads',
'proportion_PV_beads','proportion_SC_and_PV_beads','A_times_I_per_bead_forSC_alone','A_times_I_per_bead_forS
C_inPV','A_times_I_per_bead_forPV_alone','A_times_I_per_bead_forPV_inSC'};
headers=headers';
Summary=horzcat(headers,values);
Summary=cell2table(Summary,'VariableNames',{'Summarized_Parameters' 'values'});

%% save data in the folder

save(fullfile(Folder,'Particle.mat'),'TableGFP', 'TableCherry')
writetable(TableGFP, fullfile(Folder,'ParticleGFP.csv'))
writetable(TableCherry, fullfile(Folder,'ParticleCherry.csv'))
writetable(Summary, fullfile(Folder,'Summary.csv'))

%% preparing input for subsequent analysis, August 2017, By NH
% here we want to prepare an input (PV, SC, # of particle) for each recorded bead.

input_NN=zeros(a,3);

for   i=1:a
   if  Count_Total_NrBeads(i,1) < Count_Total_NrBeads(i+1,1) && list_for_Area_times_Int_PV(i)~=0
      input_NN(i,1)=list_for_Area_times_Int_PV(i);
      input_NN(i,2)=0;
      input_NN(i,3)=ParticleNr(i);

   elseif Count_Total_NrBeads(i,1) < Count_Total_NrBeads(i+1,1) && list_AtimesI_SC_only(i)~=0
      input_NN(i,1)=0;
      input_NN(i,2)=list_AtimesI_SC_only(i);
      input_NN(i,3)=ParticleNr(i);

   elseif Count_Total_NrBeads(i,1) < Count_Total_NrBeads(i+1,1) && list_AtimesI_SC_only(i)==0 &&
list_for_Area_times_Int_PV(i)==0
      input_NN(i,1)=list_SC_A_times_I_inbeads_with_PV(i);
      input_NN(i,2)=list_for_Area_times_Int_PV_with_SC(i);
      input_NN(i,3)=ParticleNr(i);



   end
end

% Remove zero rows
input_NN( all(~input_NN,2), : ) = [];
input_NN=mat2dataset(input_NN);
input_NN=dataset2table(input_NN);
writetable(input_NN, fullfile(Folder,'input_NN.csv'));

%% preparing the inut for the interaction alalysis of beeds with 2 SC particles. Calculating their surface and their
distance

SC_interaction=zeros(a,3);
```

```
for i=2:a
   if beadsWithSConly(i)==1 && ParticleNr(i)==2
       SC_interaction(i,1)= SytoArea(i,1)*SytoIntensity(i,1);
       SC_interaction(i,2)= SytoArea(i-1,1)*SytoIntensity(i-1,1);
       SC_interaction(i,3)= sqrt((table2array(TableGFP_2(i,3))-table2array(TableGFP_2(i-1,3)))^2+
(table2array(TableGFP_2(i,4))-table2array(TableGFP_2(i-1,4)))^2);
   end
end

% Remove zero rows
SC_interaction( all(~SC_interaction,2), : ) = [];
SC_interaction=mat2dataset(SC_interaction);
SC_interaction=dataset2table(SC_interaction);
writetable(SC_interaction, fullfile(Folder,'SC_interaction.csv'));

clearvars -except Main Timepoints Experiment Timepoint
end
end

%%PART2

%NH\July 2017:Accessing the input_NN file from all the folders for
%subsequent analysis. The "AllinputNN" structure in the workspace must be
%manually saved.
Main='/folderX/folderY/ExperimentFolder/';

%fixed parameters
Timepoints=dir([Main,'T*']);
delimiter = ',';
startRow = 2;
formatSpec = '%f%f%f%[^\n\r]';

%inside each time point
for t=1:length(Timepoints)
Timepoint=fullfile(Main, Timepoints(t).name);
Experiment=dir(Timepoint);
Experiment = Experiment(~strncmpi('.', {Experiment.name}, 1));

%inside each experiment
for e=1:length(Experiment)
filename =fullfile(Timepoint, Experiment(e).name,'/','input_NN.csv');

fileID = fopen(filename,'r');

%% Read columns of data according to format string.
% This call is based on the structure of the file used to generate this
% code. If an error occurs for a different file, try regenerating the code
% from the Import Tool.
dataArray = textscan(fileID, formatSpec, 'Delimiter', delimiter, 'HeaderLines' ,startRow-1, 'ReturnOnError', false);

%% Close the text file.
fclose(fileID);
file_name=strcat(Timepoints(t).name,'_',Experiment(e).name);
ALLinputNN.(file_name)= [dataArray{1:end-1}];
```

```
end
end

%% PART 3

%NH\July 2017:Accessing the SC_Interaction from all the folders for
%subsequent analysis. The "ALLSCInteraction" structure in the workspace must be
%manually saved.
Main='/folderX/folderY/ExperimentFolder/';
%fixed parameters
Timepoints=dir([Main,'T*']);
delimiter = ',';
startRow = 2;
formatSpec = '%f%f%f%[^/n/r]';

%inside each time point
for t=1:length(Timepoints)
Timepoint=fullfile(Main, Timepoints(t).name);
Experiment=dir(Timepoint);
Experiment = Experiment(~strncmpi('.', {Experiment.name}, 1));

%inside each experiment
for e=1:length(Experiment)
filename =fullfile(Timepoint, Experiment(e).name,'/','SC_interaction.csv');

fileID = fopen(filename,'r');

%% Read columns of data according to format string.
% This call is based on the structure of the file used to generate this
% code. If an error occurs for a different file, try regenerating the code
% from the Import Tool.
dataArray = textscan(fileID, formatSpec, 'Delimiter', delimiter, 'HeaderLines' ,startRow-1, 'ReturnOnError', false);

%% Close the text file.
fclose(fileID);
file_name=strcat(Timepoints(t).name,'_',Experiment(e).name);
ALLSCInteraction.(file_name)= [dataArray{1:end-1}];

end
end

%% PART 4


%NH\July 2017:Accessing the Summary file from all the folders for
%subsequent analysis. The "AllSummary" structure in the workspace must be
%manually saved.
Main='/folderX/folderY/ExperimentFolder/';

%fixed parameters
Timepoints=dir([Main,'T*']);
delimiter = ',';
startRow = 2;
```

```
formatSpec = '%*s%f%[^\n\r]';

%inside each time point
for t=1:length(Timepoints)
Timepoint=fullfile(Main, Timepoints(t).name);
Experiment=dir(Timepoint);
Experiment = Experiment(~strncmpi('.', {Experiment.name}, 1));

%inside each experiment
for e=1:length(Experiment)
filename =fullfile(Timepoint, Experiment(e).name,'/','Summary.csv');
% Open the text file.
fileID = fopen(filename,'r');

% Read columns of data according to format string.
% This call is based on the structure of the file used to generate this
% code. If an error occurs for a different file, try regenerating the code
% from the Import Tool.
dataArray = textscan(fileID, formatSpec, 'Delimiter', delimiter, 'HeaderLines' ,startRow-1, 'ReturnOnError', false);

% Close the text file.
fclose(fileID);


%% Create output variable
file_name=strcat(Timepoints(t).name,'_',Experiment(e).name);
AllSummary.(file_name)= [dataArray{1:end-1}];
% AllSummary.T6_SC_Rep1(isnan(AllSummary.T6_SC_Rep1))=0
% for m=1:36(AllSummary);
%     m(isnan(m))=0;
% end
end
end
%% Clear temporary variables
clearvars filename delimiter startRow formatSpec fileID dataArray ans;

% %--------To convert all summary into csv---------
% Alldata = struct2table(AllSummary);
% writetable(Alldata,'/filenameX/filenameY/Data.csv')
```
